# *In-cell* structure of a LINC complex reveals the molecular basis for membrane remodelling and head-to-tail coupling in sperm cells

**DOI:** 10.1101/2025.08.04.668116

**Authors:** Tom Dendooven, Mina Ebrahimi, Alia dos Santos, Warre Dhondt, Oliver Knowles, Thomas Hale, Alister Burt, Ahmad Reza Mehdipour, Matteo Allegretti

## Abstract

In eukaryotic cells, LINC complexes physically bridge the two nuclear membranes to cytoskeletal filaments, transmitting mechanical forces to the nucleus. Due to their dynamic and membrane-embedded nature, their architecture in native membranes has been hitherto elusive. Here, we combine *in-situ* electron cryo-tomography, AlphaFold predictions and molecular dynamics simulations to determine the sub-nanometer-resolution architecture of the trimeric ∼130 kDa SUN5-Nesprin3 LINC complex in human spermatozoa. The structure reveals a hexagonal lattice that stitches the nuclear envelope, creating a rigid interface that links the sperm nucleus to the flagellum. Our structure unveils a unique membrane-anchoring mechanism: a SUN5 *KASH-lid* β-hairpin and a Nesprin3 amphipathic helix insert into the outer nuclear membrane, leading to cooperative remodelling of the nuclear envelope shape by the extended LINC lattice, which results in flattening the caudal sperm nucleus. Our integrative data provides key insights into the architecture and role of a LINC complex *in situ*, revealing molecular details of a severe male infertility phenotype.

## Introduction

LINC (**LI**nkers of **N**ucleoskeleton and **C**ytoskeleton) complexes are fundamental for several cellular processes, including nuclear positioning and reshaping, cell polarization and migration, chromosome pairing, and mechano-regulation of gene expression (Friedl *et al*, 2011; J. David Pajerowski, 2007; Lombardi *et al*, 2011; Rothballer & Kutay, 2013; Starr & Fridolfsson, 2010; Wang, 2009). These assemblies typically consist of homotrimeric SUN proteins (Sad1p, UNC-84 homology domain-containing proteins) (Hennen *et al*, 2018; Lu *et al*, 2008; Zhou *et al*, 2012) bound to KASH proteins (Klarsicht, ANC-1, Syne1 Homology domain-containing proteins), forming a supra-molecular assembly in the perinuclear space (McGillivary *et al*, 2023; Sosa *et al*, 2013). SUN proteins comprise an N-terminus interacting with chromatin domains and lamins in metazoans, a transmembrane segment, a coiled-coil region, and a C-terminal SUN domain. KASH proteins interact with the SUN domain through a conserved C-terminal KASH domain (Sosa *et al*, 2012), and contain an N-terminal cytoplasmic domain that binds all major cytoskeletal filaments (McGillivary *et al*., 2023). LINC complexes also play essential roles in sperm differentiation (Gob *et al*, 2010), where they direct meiotic chromosome movement, acrosome formation and nuclear remodelling (Cavazza *et al*, 2021; Chaigne *et al*, 2016; Donahue, 1972; Kmonickova *et al*, 2020; Scheffler *et al*, 2021).

Although LINC complexes play crucial roles in numerous cellular functions, their native structure and oligomeric state at the nuclear membrane remain elusive.

In this study, we investigate the native architecture of SUN5 LINC complexes that are part of the Head-to-Tail Coupling Apparatus (HTCA) in mature human spermatozoa, where they connect the inner (INM) and the outer (ONM) sperm nuclear membranes, transmitting mechanical forces from the flagellar cytoskeleton (Galletta *et al*, 2020; Rothballer & Kutay, 2013; Wu *et al*, 2020). SUN5 is a sperm-specific SUN-domain protein expressed at the spermatocyte differentiation stage (Xian-Zhen Jiang, 2011) that interacts with the KASH protein Nesprin3 to form a functional LINC complex (Manfrevola *et al*, 2021; Morgan *et al*, 2011; Zhang *et al*, 2021b). *Sun5* is the most frequently implicated gene in patients with Acephalic Spermatozoa Syndrome (ASS), with biallelic loss-of-function variants identified in approximately 33–47% of ASS patients across diverse populations (Zhu et al., 2016; Sha et al., 2018b; Göb et al., 2010; Shang et al., 2017(Xiang *et al*, 2022). ASS is a severe form of teratozoospermia where affected men present nearly 100% headless (or tailless) sperm cells and suffer from severe male-factor infertility due to a failure in the HTCA assembly (Galletta et al., 2020; Wu et al., 2020; Avidor-Reiss et al., 2020). In addition, ultrastructural studies of spermatogenesis in *Sun5*^-/-^ male mice revealed that, while the sperm HTCA can be successfully assembled during the early stage of spermiogenesis, it disassociates during spermatid elongation, resulting in acephalic mature spermatozoa (Shang *et al*, 2017).

Using *in situ* cryo-electron tomography (cryo-ET) and sub-tomogram averaging, we resolved the native layout of the SUN5-Nesprin3 LINC complex spanning the basal plate of human motile spermatozoa at sub-nanometer resolution. Our reconstruction reveals that the SUN5-Nesprin3 LINC complex assembles into a hexagonal lattice embedded at the sperm HTCA. Through a combination of AlphaFold-guided density fitting and atomistic molecular dynamics (MD) simulations, we describe the molecular details of how a SUN5 KASH-lid forms a membrane anchor that embeds in the sperm outer nuclear envelope (NE) in a Nesprin3-dependent manner. We map SUN5 mutations recorded in ASS patients onto our structure, rationalising their structural contribution to the acephalic sperm phenotype. Finally, we use atomistic MD to propose a SUN5-Nesprin3 lattice-driven mechanism of NE curvature inversion, which has physiological importance in forming the *implantation fossa* depression at the caudal side of the sperm nucleus. Overall, our integrative structural biology approach provides key insights into the higher-order molecular assembly of a membrane-bound LINC complex *in situ*, its contribution to shaping the nuclear membrane, and the molecular basis underlying a severe male-factor infertility phenotype.

## Results

### A 2D hexagonal lattice of SUN5 trimers spans the basal plate of the *implantation fossa*

We used cryo-FIB milling to thin vitrified sperm cells for *in-situ* cryo-ET studies of the sperm neck region. We collected ∼230 tilt-series (Eisenstein *et al*, 2023) of the *implantation fossa* (Fig. 1A) and our reconstructed tomograms revealed small protein assemblies spanning the INM and ONM of the sperm basal plate region in a regular pattern (Fig. 1B). Through subtomogram averaging (Burt *et al*, 2021), we resolved a 9.4 Å resolution reconstruction of a two-dimensional hexagonal lattice sitting on the perinuclear side of the ONM (Fig. 1C, Fig. S1A-C). The resulting map showed that a defined 12nm gap between the INM and ONM (Fig. 1C), approximately half of the canonical INM-ONM distance in somatic cells (Kucinska *et al*, 2023), is decorated by trimeric clover-like assemblies (Fig. 1C-D). From these trimeric assemblies, density for a triple coiled coil of ∼4.5nm length emanates, connecting the clover structures to the INM (Fig. 1C). These features are characteristic of SUN protein trimers (Zhou *et al*., 2012), where the C-terminal SUN domain folds into a β-sandwich and is preceded by a triple-coiled coil element. To map the correct SUN protein to our sub-tomogram average, we combined immunostaining with AlphaFold3 structure prediction (Abramson *et al*, 2024) and density fitting in the sub-tomogram average. In agreement with observations from previous studies, we found that SUN5 is precisely localized at the basal plate of the HTCA (Fig. 1B), in-between the redundant NE membranes containing nuclear pore complexes (Cazin *et al*, 2021; Santos *et al*, 2024; Shang *et al*, 2018; Zhang *et al*, 2021a; Zhu *et al*, 2018) (Fig. S2). We further showed that SUN5 is expressed at the spermatocyte stage of sperm differentiation, after which it accumulates at the nuclear posterior (the *implantation fossa)* (Fig. S3). AlphaFold3 predictions of SUN5 trimers, containing both the SUN β-sandwich domain and the coiled coil region, fitted well into our map (Fig. 1D-E, Fig. S4A). Notably, the SUN5 triple-coiled coil is shorter than for other trimeric SUN proteins, as previously hypothesised based on the SUN5 sequence, explaining the short 12nm distance between the INM and ONM observed in our tomograms (Fig. 1C, Fig. S4B) (Jahed *et al*, 2016; Shang *et al*., 2017; Sosa *et al*., 2013). The interface between SUN domains of neighbouring SUN5 trimers in the hexagonal lattice is mainly mediated by hydrophobic, Van der Waals, and electrostatic interactions, as has been previously postulated *in silico* for SUN1 (Jahed *et al*, 2018) (Fig. S4D). In summary, we solved the architecture of the ∼130kDa SUN5 trimer *in situ* at subnanometer resolution and show that it organises into a hexagonal 2D lattice at the NE (Fig. 1C-E, Fig. S1A, Movie Supp. 1), forming the basal plate at the caudal side of the sperm head.

**Fig. 1.**
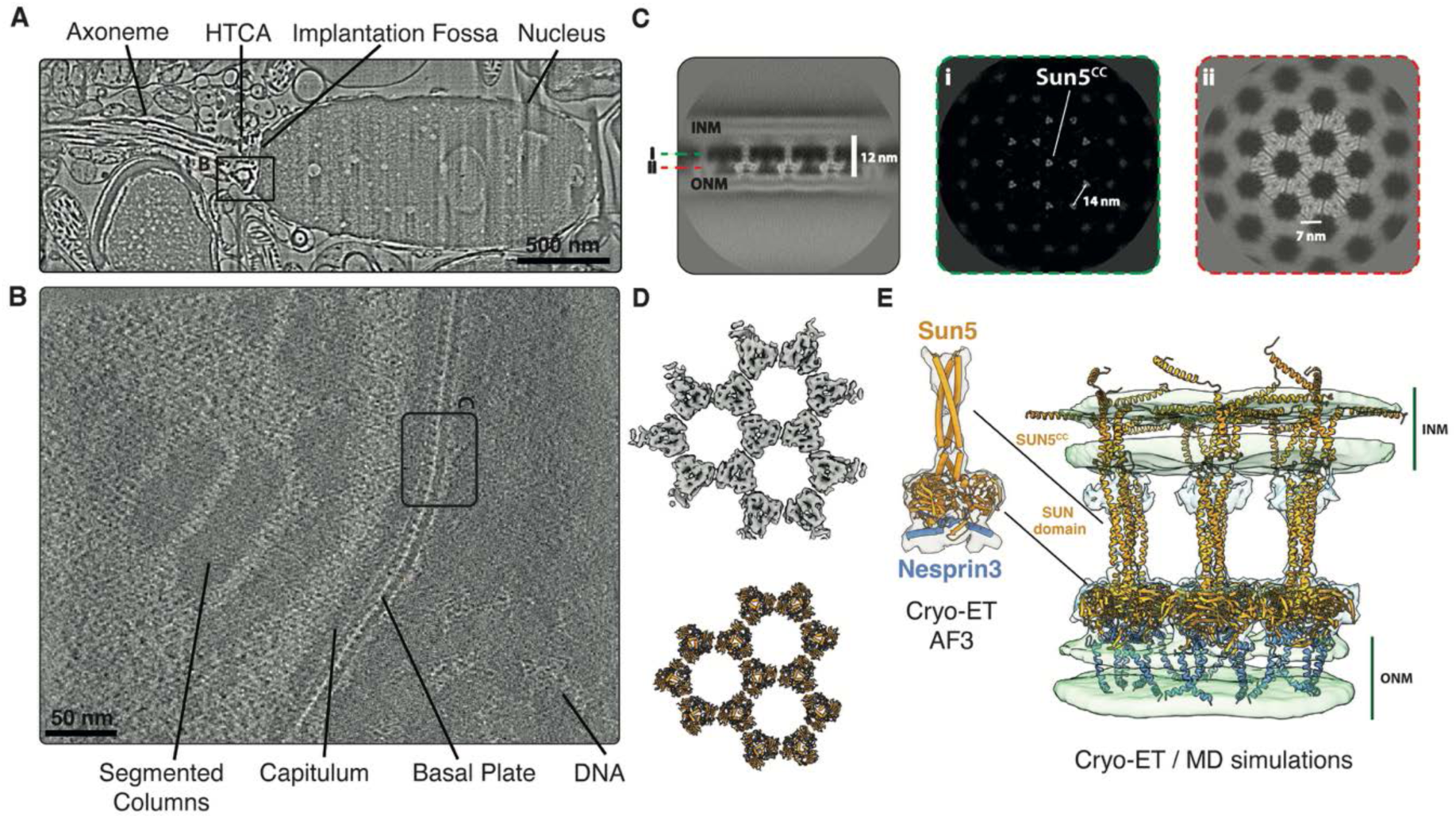
The *in-cell* architecture of the SUN5-Nesprin3 complex at the HTCA. A) Cryo-SEM image of a vitrified sperm cell, with the axoneme, *head-to-tail* coupling apparatus (HTCA), *implantation fossa* and nucleus annotated. The rectangular inset is described in B). **B**) Tomographic slice through a high-magnification tomogram of the sperm neck, showing the segmented columns, the capitulum, DNA and the basal plate, as well as a part of the NE. A regular protein pattern spans the NE. The rectangular inset is described in C. **C**) Slices through the sub-tomogram average of the basal plate, revealing a hexagonal lattice of homo-trimeric SUN5-Nesprin3 LINC complexes. A triple coiled-coil fills the 12nm gap between the INM and ONM of the basal plate, formed through lattice contacts between trimers of the SUN5-domains. **D**) Surface map and model of the *in-situ* hexagonal lattice decorating the basal plate. **E**) Docked model of the SUN5-Nesprin3 trimers in the sub-tomogram average, refined by molecular dynamics-based density fitting.

### Molecular dynamics simulations reveal Nesprin3-mediated exposure of a SUN5 membrane anchor for ONM binding

SUN5 was previously shown to bind the KASH protein Nesprin3 (Manfrevola *et al*., 2021; Morgan *et al*., 2011; Zhang *et al*., 2021b). Our immunostaining data confirmed this, with Nesprin3 signal localising at the nuclear posterior, together with SUN5 (Fig. S2). Indeed, AlphaFold3 predictions of SUN5 bound to the C-terminal KASH domain of Nesprin3 revealed an exposed β-hairpin KASH-lid at the base of the SUN domain of SUN5 (Fig. S4C), in agreement with our sub-tomogram average (Fig. 1E), and as was observed in a SUN2-Nesprin3 crystal structure and several other SUN-KASH *in vitro* structures (Cruz *et al*, 2020; Gurusaran & Davies, 2021; Gurusaran *et al*, 2024; Sosa *et al*., 2012; Wang *et al*, 2012; Zhou *et al*., 2012). To obtain a more accurate model of how the SUN5-Nesprin3 lattice interacts with the INM and ONM, we carried out atomistic MD simulations using our subtomogram averaging-derived model as a starting point (Fig. 1E, Fig. 2A, Fig. S5). Interestingly, AlphaFold3 predictions showed that the KASH-lid of SUN5 folds into a structured β-hairpin only when the C-terminal KASH domain of Nesprin3 is bound (Fig. S4C), in agreement with crystal structures of other SUN paralogs (Cruz *et al*, 2020; Sosa *et al*., 2012). While the KASH-lid sequence is not strongly conserved among SUN proteins, it is nevertheless rich in hydrophobic and aromatic residues in all five SUN paralogs in humans (Sosa *et al*., 2012; Wang *et al*, 2012). Notably, Trp232 is highly conserved in SUN protein orthologs, while Trp226 is conserved in SUN3 and SUN4 paralogs (Sosa *et al*., 2012). We performed atomistic MD of SUN5 anchoring in both solution and in the ONM lipid bilayer in the presence or absence of the Nesprin3 KASH domain (Fig. 2B, Fig. S6C-D). The SUN5 KASH-lid inserts into the lipid bilayer via a “Trp-anchor” motif, composed of Trp226, Trp228 (unique to SUN5), and Trp232 (Fig. 2A). This finding is consistent with *in-cell* studies showing impaired membrane binding upon KASH-lid deletion in SUN2 (Sosa *et al*., 2012). Notably, our simulations reveal that Nesprin3 binding is crucial for maintaining the stable membrane association of SUN5 (Fig. 2B, Fig. S6C-D), in agreement with our AlphaFold3 predictions (Fig. S4, Movie Supp. 2). During the simulations of the SUN domain in solution, the KASH-lid β-hairpin becomes unstable in the absence of Nesprin3 (Fig. S6A-B). However, in simulations with the ONM, we observed a form of molecular hysteresis: while Nesprin3 is required for the formation of the stable KASH-lid β-hairpin, once the hairpin is formed and anchored into the ONM, it remains stable even after Nesprin3 is removed *in silico* (Fig. S6C-D). In summary, our data reveal that the Nesprin3 KASH domain acts as a molecular chaperone, inducing the folding and exposure of the SUN5 KASH-lid β-hairpin for tight anchoring of the SUN5 LINC complex to the ONM (Fig. 2A-B, Fig. S6).

**Fig. 2.**
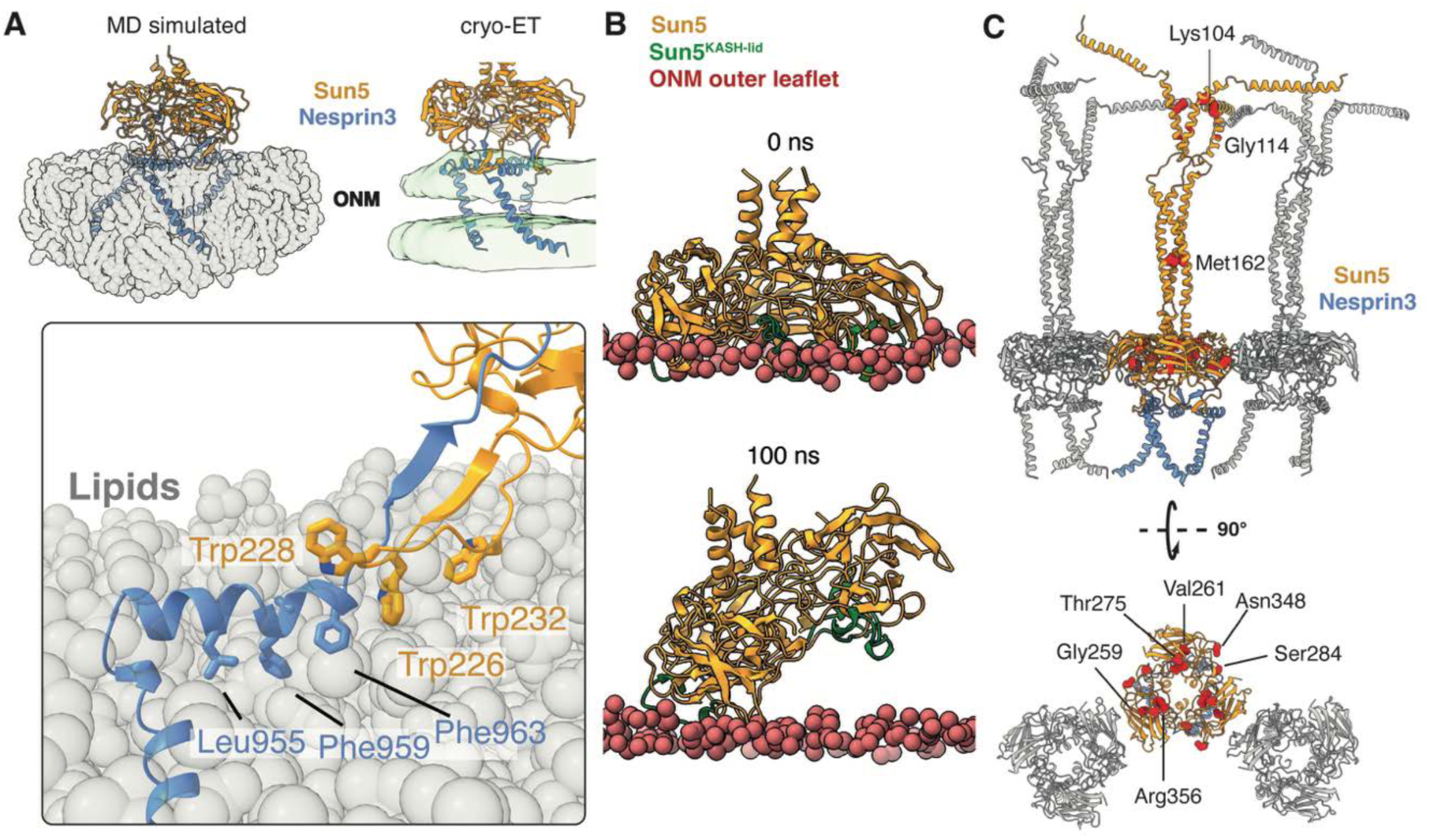
Membrane anchorage and infertility mutations. **A**) MD simulation of the SUN5-domain interaction with the ONM. Three Trp residues (226, 228, and 232) of the SUN5 KASH-lid form a membrane anchor, in addition to a Nesprin3 amphipathic helix N-terminally of the Nesprin3 KASH domain. A Nesprin3 transmembrane helix extends from the amphipathic helix to traverse the ONM. **B**) MD simulation of the SUN5-domain interaction in the absence of Nesprin3 and with the KASH-lid folded into the SUN domain, where SUN5 dissociates from the membrane on a sub-microsecond timescale. **C**) Structural mapping of SUN5 residues known to be mutated in patients with Acephalic Sperm Syndrome (ASS). Mutations are predicted to interfere with SUN5 domain folding and trimerisation, hexagonal lattice formation, SUN5 triple coiled coil folding and INM docking.

N-terminally of the KASH domain of Nesprin3, a conserved amphipathic helix (aa: 950-963), lies flat on the ONM inner leaflet in our simulations (Fig. 1E, Fig. 2A) (Sosa *et al*., 2012). Lastly, a Nesprin3 transmembrane helix (aa:919-946) then traverses the ONM bilayer, positioning the Nesprin3 spectrin repeat domain to the cytoplasmic side of the NE and likely embedded in the sperm capitulum (Zhang *et al*., 2021b) (Fig. 1B, Fig. 2A).

Notably, several SUN5 mutations have been found in patients with severe male-factor infertility (Sha *et al*, 2018; Xiang *et al*., 2022; Zhu *et al*, 2016). These mutations result in a defective HTCA and acephalic spermatozoa in the ejaculate, while the severed sperm heads remain in the seminiferous epithelium of the testis (Zhu *et al*., 2016). These mutations have been shown to impair the localisation and integration of SUN5 into the NE (Shang *et al*., 2018), thereby precluding the formation of the higher-order LINC lattice resolved in this study. We mapped all known SUN5 mutations leading to ASS syndrome in the context of our SUN5-Nesprin3 lattice model (Fig. 2C). Several mutations were in critical regions of SUN domain folding, SUN5 trimerization, binding of the Nesprin3 KASH-domain, coiled-coil formation or INM binding (Gly259Ser, Val261Met, Thr275Met, Ser284Ala, Asn348Iso, Arg356Cys, Gly114Arg) (Shang *et al*., 2018).

### The SUN5-Nesprin3 lattice reverses membrane curvature at the *implantation fossa*

In addition to the significant reduction in INM-ONM distance (12nm) at the basal plate by the SUN5-Nesprin3 lattice, our cryo-TEM and SEM tomograms show a strong flattening of the nuclear membrane lining the capitulum, generating the *implantation fossa*’s depression at the sperm head posterior (Fig. 1A-B). We used atomistic MD simulations to explore whether the SUN5-Nesprin3 lattice contributes to this drastic remodelling of the nuclear membrane. We simulated different SUN5-Nesprin3 oligomers embedded in the ONM with increasing complexity, ranging from the trimeric form (Fig. S7A, Fig. S5), to a hexamer of trimers (Fig. S3A, Fig. S8, Fig. S5), a trimer of hexamers (Fig. S8, Fig. S5), and a ring of hexamers around a central hexamer (minimal lattice, seven hexamers in total) (Fig. S8, Fig. S5). We also simulated the hexameric state embedded in both the INM and ONM (Fig. S7B, Fig. S5). A single SUN5-Nesprin3 trimer induced local ONM distortion/bending of the membrane at the KASH-lid-membrane interface, in good agreement with our sub-tomogram average (Fig. S7A). In contrast, the hexameric SUN5-Nesprin3 ring, trimer of hexamers, the minimal lattice, and the double membrane system all generated a significant global curvature of the ONM (Fig. 3, Fig. S7B-C, Fig. S8). Next, we explored the contribution of Nesprin3 domains to membrane bending by simulating the hexameric form in three different states: SUN5 alone, SUN5 with the KASH domain of Nesprin3 bound, including the amphipathic helix (aa:948:975), and SUN5 bound to the KASH domain of Nesprin3, including the amphipathic helix and the transmembrane helix (aa:901-975) (Fig. 3A). Interestingly, a hexamer of SUN5 alone did not induce significant curvature. Binding of the Nesprin3 KASH domain and amphipathic helix, however, was sufficient to induce membrane curvature (Fig. 3A). Inclusion of the transmembrane helix did not increase membrane bending further, which is consistent with the observed trend in ER proteins (Bhaskara *et al*, 2019) (Fig 3A). We then replaced the KASH domain (including the amphipathic helix) with the Nesprin2 sequence in our simulations (SUN5-Nesprin2) and observed that the presence of Nesprin2 can also induce membrane curvature to a similar extent as Nesprin3 (Fig. S10). To quantify the NE membrane remodelling by a continuous SUN5-Nesprin3 lattice, we measured the curvature inversion of the basal plate in cryo-Scanning Electron Microscopy (SEM) tomograms of vitrified whole sperm cells (Fig. 1A, Fig. 3B). This region exhibited a distinctly flattened membrane profile compared to the adjacent areas. We quantified this effect by first estimating the expected curvature of an idealized, unperturbed membrane by fitting a spline to the two adjacent, non-flattened regions (Fig. 3B). By comparing the differential curvature between this ideal curve and the observed flattened area, we calculated the magnitude of the SUN5-induced straightening to be approximately 1.75±0.23 ¼m^−1^ (Table S1). We then tested the hypothesis that the formation of the highly ordered SUN5-Nesprin3 lattice is the primary driving force for the membrane straightening. We connected our experimental observations with the simulated molecular-scale behaviour by comparing the measured 2D curvature from SEM tomograms to the principal curvature calculated from simulations of a single SUN5-Nesprin3 hexamer, a trimer of hexamers, and a minimal lattice. Interestingly, plotting the principal curvature (k_1_, Fig. 3B) from these simulations against the length of the simulated SUN5-Nesprin3 complex revealed a power-law relationship, which appears linear on a log-log plot (Fig. 3B). The resulting regression provides a good predictive model that directly links the size of a SUN5-Nesprin3 lattice to the magnitude of the membrane curvature remodelling it induces.

**Fig. 3.**
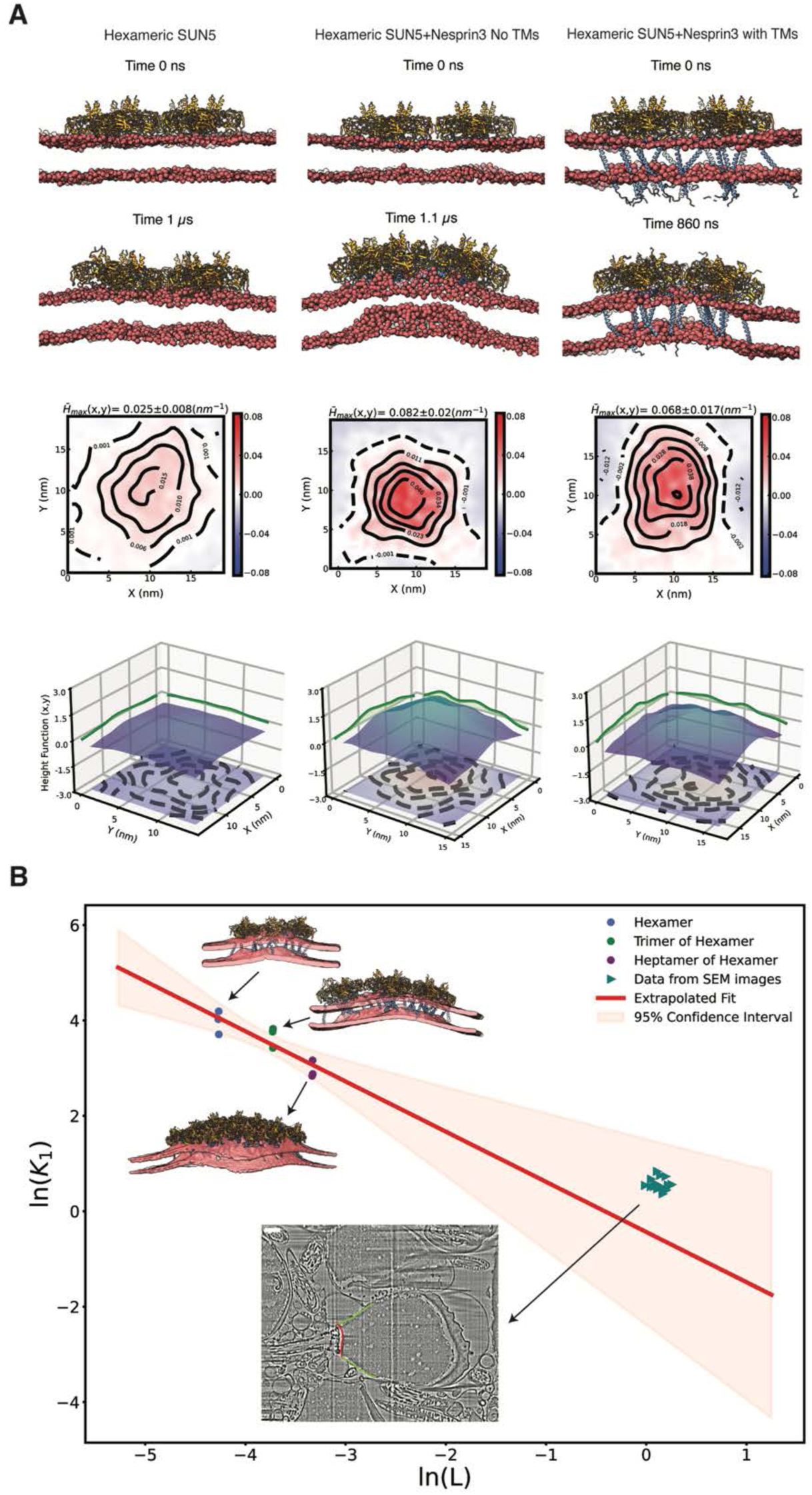
The *curvature-inducing* property of the SUN5-Nesprin3 higher-order complexes. **A**) SUN5 hexamer (left) does not perturb the bilayer significantly, while SUN5-Nesprin3 (without transmembrane helices) (middle) and SUN5-Nesprin3 (with transmembrane helices) (right) hexamers strongly perturb the bilayer structure. Side view snapshots at the beginning of the simulations (top panels) and the end of the simulations (upper middle panels) with SUN5 coloured orange, Nesprin3 coloured blue, and Membrane (P atoms) coloured red. TM stands for transmembrane helices. Lower middle panels show contour maps of the averaged curvature profiles for each system, highlighting the extent of curvature induction around the centre of the SUN5-Nepsrin3 hexamer complex. The maximum value of curvature fields and Std. is shown at the Top. Bottom panels show local membrane shape approximated by a height function of the midplane (height contour map along the plane). **B**) Power-law fit of the log-transformed principal curvature (k1) versus the radius of the complex from simulations of a hexamer, a trimer of hexamers, and heptamers of hexamers. The solid red line represents the fit within the interpolated and the extrapolated regions. The shaded area is the 95% confidence interval for the fit. The green triangles represent the 2D flattening curvature calculated from the SEM images (not used in the fit). In the SEM panel image, the red and green lines show the tracing of the SUN5-containing flattened area and the two adjacent membranes, respectively (see Materials and Methods and Table S1 for details).

To further corroborate these results *in-cell*, we overexpressed SUN5 in human fibroblasts, which endogenously express Nesprin3 (Fig. S9F). After cryo-FIB milling and tilt-series acquisition, we carried out a morphometric analysis of the NE (Barad *et al*, 2023) and compared it with the NE of wild-type fibroblasts. The nuclear membrane in the SUN5 overexpressed cells frequently showed invaginated areas with high curvature (Fig. S9A, C), observed in 65% of the tomograms analysed, which were absent in wild-type cells (Fig. S9A, D). This increased the overall nuclear membrane curvature (Fig. S9B). Notably, the INM-ONM distance was very heterogeneous upon overexpression of SUN5, with two prominent peaks in the histogram, in comparison to control cells (Fig. S9C-E). One peak was at the expected perinuclear space distance for human fibroblasts (around 22nm) while another peak was at around 10nm, close to the INM-ONM distance observed in the sperm basal plate (Fig. 1B-C, Fig. S9C). Overall, the *in situ* and *in silico* analyses of the SUN5-Nesprin3 lattice reveal a striking capacity for NE remodelling, which shapes the mature sperm head.

## Discussion

In this study, we determined the *in-situ* structure of the SUN5-Nesprin3 LINC complex, at sub-nanometer resolution in motile human spermatozoa using sub-tomogram averaging. Our AlphaFold3 predictions and atomistic MD simulations also describe the mechanistic basis for how LINC architecture is connected to NE remodelling and male fertility. Studies of LINC complexes in their native, double lipid membrane-embedded state have been intractable due to the lack of suitable *in vitro* reconstitution systems with two membrane bilayers. In fact, all available structures solved by X-ray crystallography contained only the solubilised C-terminal SUN domains with or without KASH-domain peptides bound (Cruz *et al*., 2020; Gurusaran & Davies, 2021; Gurusaran *et al*, 2024; Sosa *et al*., 2012; Wang *et al*., 2012; Zhou *et al*., 2012), drawbacks that we overcame with our *in situ* cryo-ET approach. Our reconstruction of the native SUN5 LINC complex revealed a hexameric lattice of SUN5-Nesprin3 particles (Fig. 1). SUN5 is an essential protein for spermatogenesis (Shang *et al*., 2017), and SUN5 mutations are implicated in approximately 40% of ASS cases (Cazin *et al*., 2021; Shang *et al*., 2017; Xiang *et al*., 2022; Zhu *et al*., 2016). As a result, an in-depth structural characterisation was crucial to understand the molecular basis of SUN5-related male infertility. The higher-order hexagonal architecture we have determined is mediated exclusively by lateral interactions of neighbouring SUN5 trimers’ β-sandwiches (Fig. 1C-E, Fig. S4D) (Jahed *et al*., 2018). A hexameric organisation of SUN domains *in situ* has been previously hypothesised for SUN1 complexes, based on MD simulations and fluorescence fluctuation spectroscopy measurements (Hennen *et al*., 2018; Jahed *et al*., 2018; Wang *et al*., 2012; Yerima *et al*, 2023; Zhou *et al*., 2012). Recently, other models for SUN-KASH high-order assemblies have been proposed and experimentally validated *in vitro,* where SUN1 trimers interact head-to-head via KASH-lids or KASH peptides (Gurusaran & Davies, 2021; Gurusaran *et al*., 2024), although, in light of our findings, the physiological relevance of these remains to be proven (McGillivary et al., 2023). The KASH partner of SUN5 is Nesprin3, as determined by our immunostaining (Fig. S2) and in agreement with previous work (Manfrevola *et al*., 2021; Morgan *et al*., 2011; Zhang *et al*., 2021b). It was previously shown that Nesprin3 plays a role in centriole attachment to the NE as part of the HTCA, and a knockdown of Nesprin3 increases the distance between the nucleus and the centriole (Morgan *et al*., 2011).

A central insight from our work is the role of membrane-anchored protein elements in driving NE remodelling. The SUN5 *KASH-lid* – a β-hairpin at the base of the SUN domain – contains an unusually extended tri-tryptophan motif unique to SUN5 (Fig. 2A) (Sosa *et al*., 2012). We find that this tryptophan-rich hairpin acts as a reversible membrane anchor on the inner leaflet of the outer nuclear membrane (ONM). Tryptophan and other aromatic residues have well-known interfacial “anchor” properties in lipid bilayers, inserting into the hydrophobic/hydrophilic interface to stabilise proteins at membranes (Karl Salzwedel, 1999; Kundu *et al*, 2023; Sanchez *et al*, 2011). Our atomistic simulations showed that in SUN5, the KASH-lid remains unstructured and disengaged in the absence of a KASH partner, but upon binding Nesprin3 it folds into the β-hairpin and *wedges* into the ONM (Fig. 2B, Fig. S6C-D). Thus, Nesprin3 binding “activates” the SUN5 KASH-lid as a membrane anchor, mechanically coupling the inner and outer nuclear membranes. This mechanism creates a tight LINC linkage across the perinuclear space without requiring SUN5 to have a classical transmembrane extension into the ONM.

From our cryo-tomography data and simulations, membrane insertion by the SUN5 KASH-lid and the Nesprin3 amphipathic helix emerges as the driver of NE curvature change. In our simulations, hexameric SUN5 alone did not significantly deform a lipid bilayer (Fig. 3A). However, when SUN5 is bound to a Nesprin3 KASH domain including the short amphipathic helix, pronounced membrane bending occurs for higher-order LINC assemblies (Fig. 3B, Fig. S7A-B, Fig. S8). Notably, the concept of a partially embedded membrane anchor as a curvature modulator is not unique to SUN5. ER-shaping proteins like the reticulon-homology protein FAM134B use pairs of intramembrane helices and amphipathic wedges to bend ER membranes (Bhaskara *et al*., 2019), and even nuclear pore complexes rely on amphipathic motifs (ALPS helices) in certain nucleoporins to stabilize the curved pore membrane (Meszaros *et al*, 2015). Our findings extend this paradigm to the sperm NE, with the SUN5 KASH-lid representing a specialised, membrane-anchored “wedge” that is deployed during sperm nuclear remodelling. By quantitatively measuring the sperm basal plate’s curvature and comparing it adjacent nuclear regions in SEM cryo-tomograms, we showed that the LINC lattice straightens the NE curvature consistent with our multiscale simulations (Fig. 1A-C, Fig. 3B). Interestingly, the Nesprin3 transmembrane helix does not play a crucial role in membrane bending, and may only serve as a connector to the sperm cytoplasm (Fig. 3A, Fig. S7), analogous to the FAM134B protein in the endoplasmic reticulum (Bhaskara *et al*., 2019). We propose that the SUN5– Nesprin3 lattice acts as a nanoscale “*molecular vise*” clamping the nuclear membranes together at the sperm basal plate and remodelling its curvature to create the *implantation fossa* and enable attachment of the sperm tail. Indeed, SUN5 deletion has been shown to cause shorter and wider sperm heads compared to wild-type ones (Shang *et al*., 2017), corroborating its proposed role in NE reshaping. Looking beyond the sperm cell, it is intriguing to consider that clustering of SUN–KASH complexes could locally modulate the curvature of the NE in response to mechanical stress or during nuclear migration. Interestingly, when we replaced Nesprin3 with Nesprin2 in our simulations, membrane bending is induced as well (Fig. S10), pointing to a more general mechanism where higher-order LINC assemblies induce membrane curvature remodelling. It is conceivable that similar SUN–KASH lattice formations, or at least small oligomeric clusters, might occur in other cell types under high mechanical load (for example, at transmembrane actin attachment sites in muscle cells or migrating fibroblasts). These speculative extensions will require further *in situ* studies of native LINC complexes. As a proof of principle, we overexpressed SUN5 in human fibroblasts (Majumder *et al*, 2022; Shang *et al*., 2018; Turgay *et al*, 2010). There, SUN5 had the chance to interact with Nesprin3 if local increases in membrane tension reduced the perinuclear space (Fig. S9C) (Zimmerli *et al*, 2021). Our data suggest that SUN5-Nesprin3 binding drives the changes in membrane curvature observed in our tomograms (Fig. S9A-C), which supports our general hypothesis. Unfortunately, in SUN5 overexpressed cells, we could not observe lattice formation, probably due to the diffuse localisation of Nesprin3 in somatic cells, or to the fact that SUN5 clustering may depend on other factors in earlier stages of sperm differentiation (Morgan *et al*., 2011). Nevertheless, our findings pave the way by demonstrating that ordered LINC arrays can exist and exert tangible effects on membrane architecture *in vivo*.

In summary, our *in-cell* cryo-EM structure of the ∼130kDa double-membrane-embedded protein SUN5, shows a high-order hexagonal assembly of a LINC complex which stitches and remodels the two nuclear membranes at the caudal side of the sperm head (Fig. 4). This structural organization is physiologically crucial to keep the sperm nucleus, which has to be delivered to the oocyte, and the tail, which drives the movement, strongly linked, as shown by the many SUN5 mutations causing severe male-factor infertility due to headless sperm in the ejaculate (Shang *et al*., 2018; Shang *et al*., 2017; Xiang *et al*., 2022; Zhu *et al*., 2016) (Fig. 2C). Our work provides a structural framework to find solutions for ASS with higher success rates than canonical ICSI (Shang *et al*., 2017), and, on the other hand, to inform the development of male contraceptives. A recent example of such a strategy is the targeting of the testis-enriched protein ARRDC5: knockout of *Arrdc5* in mice causes severe oligo-terato-asthenospermia and infertility, therefore ARRDC5 has been proposed as a promising male contraceptive target (Giassetti *et al*, 2023). SUN5 and its binding partners represent a similarly attractive target space, given their testis-specific expression and essential role in sperm assembly. By revealing how a membrane-anchored LINC complex couples organelle architecture to cell function, we not only explain a key aspect of sperm biology and male fertility but also highlight fundamental design principles by which cells use partial membrane insertion and protein self-assembly to sculpt (nuclear) membranes.

**Fig. 4.**
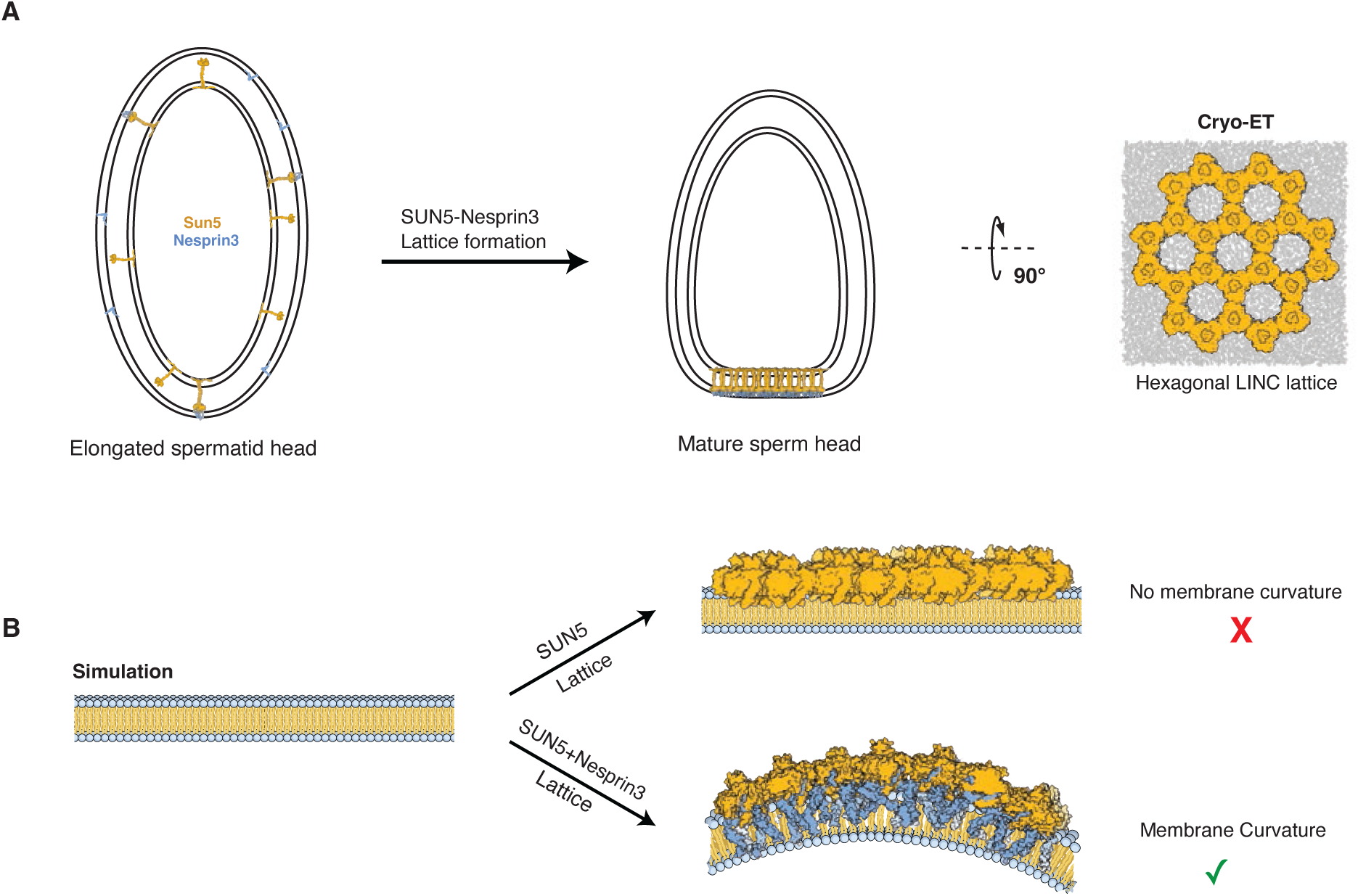
Model for remodelling of sperm nuclear envelope by SUN5-Nesprin3 lattice. **A**) a plausible process of NE curvature flattening by SUN5-Nesprin3 lattice formation. **B**) Our atomistic MD simulations results uncover that the SUN5-Nesprin3 lattice can induce positive curvature in a flat membrane, while a lattice of SUN5 alone cannot do it.

## Acknowledgements

We thank the Barford lab members for useful discussions and critical feedback. We thank Delphine Larrieu for providing hTERT human skin fibroblasts. We thank Andrew Carter, Emmanuel Derivery, Buzz Baum, Ramchandran Bhaskara, and Maziar Heidari for insightful scientific discussions. We thank the cryo-EM Facility at the LMB for maintaining the electron microscopes, the Scientific Computing facility for maintaining our computer cluster and the Light Microscopy Facility for maintaining light microscopes. We acknowledge Diamond for access and support at the Aquilos-II of the UK national electron Bio-Imaging Centre (eBIC). For the purpose of open access, the MRC Laboratory of Molecular Biology has applied a CC BY public copyright licence to any Author Accepted Manuscript version arising. The VSC (Flemish Supercomputer Center) and the EuroHPC supercomputer LUMI provided the high-performance computing resources and services used for performing the simulations in this work.

## Author Contributions

Conceptualization and experiment design: TD, ARM, MA

Writing: TD, ARM, MA, with the support of all the authors

Cryo-FIB milling and cryo-ET data collection: TD, AdS

Immunohistochemistry: AdS

Immunocytichemistry, SIM and confocal microscopy: OK, AdS

Data analysis: TD, MA analysed cryo-ET data; AB contributed with a script for data analysis;

MD simulations: ME, WD, ARM

Optical microscopy analysis: AdS

Morphometrics analysis: TH

Supervision: ARM, MA

Funding acquisition: ARM, MA, AdS

## Data and material availability

Cryo-EM map of the SUN5-Nesprin3 lattice has been deposited in the Electron Microscopy Data Bank (EMDB) with the accession code EMD-54371.

## Materials and Methods

### Human sperm and testis tissue samples

Human mature sperm cells from 25 healthy donors were obtained from the European Sperm Bank (https://www.europeanspermbank.com/en). The samples had been previously processed and tested by the European Sperm Bank according to the requirements of the national competent authorities, GMP and WHO standards, as well as EU directives: 2006/86/EC, 2006/17/EC, 2004/23/EC, 2015/565. Sperm cells were screened for morphology, genetic diseases, blood-borne viruses and motility. Consent to use the samples for research purposes was obtained by the European Sperm Bank. Human sperm straws were thawed and used immediately. Before grid vitrification or immunostaining experiments on glass, sperm cells were checked for mobility using an optical microscope.

Human testis FFPE sections (approximately 5μm thick) from five healthy donors, between 18 and 55 years of age and with no history of infertility, were obtained from the Imperial College Healthcare Tissue Bank. All the experimental procedures were approved by the Imperial College Healthcare Tissue Bank (Research Ethics Committee approval numbers: 22/WA/0214; ICHTB HTA license: 12275).

### Cell culture

Human skin fibroblasts derived from AG10803 and immortalised with SV40LT and TERT were a gift from Dr Delphine Larrieu. Cells were cultured in Dulbecco’s modified Eagle’s medium, supplemented with 10% foetal bovine serum (FBS) and penicillin/streptomycin. Cells were maintained at 37°C, 5% CO_2_.

For transient transfections, human fibroblasts were seeded on glass coverslips #1.5 or on 6-well tissue culture dishes overnight. Transfections of FLAG-SUN5 (pcDNA3.1 FLAG-SUN5 plasmid, Invitrogen) were performed using Lipofectamine 3000 (Invitrogen) for 48h. For grid preparation, non-transfected and transfected cells growing in 6-well plates were detached after 48h using 0.25% Trypsin-EDTA (Gibco) and seeded on EM grids as described in ‘*grid preparation’*.

### Grid Preparation

EM R1/4 Au 200 mesh Quantifoil grids (SPT Labtech) were glow discharged at 30mA for 45seconds on both sides, using an Edwards S150B glow discharger. Thawed sperm cells were applied directly to the grid and vitrified in liquid ethane using a Leica GP2 Plunge Freezer (Leica) or with a custom-made manual plunger. An iterative sample application and blotting prior to plunge freezing were used to assemble several sperm cell monolayers on the grid.

For human fibroblasts samples, after glow-discharging, grids were functionalized with 50μg/mL fibronectin bovine plasma (Sigma) overnight at 4℃. Human fibroblasts (WT) or fibroblasts transiently expressing Flag-SUN5 were seeded on these grids and left to adhere for 4-5 hours at 37°C, 5% CO_2_, before plunge freezing in liquid ethane using a Leica GP2 Plunge Freezer (Leica).

### Sample thinning with cryo-Focused Ion Beam milling

Vitrified cells (sperm cells and human fibroblasts) were thinned, through cryo-FIB milling, to 150-180 nm lamellae, using a Scios cryoFIB/SEM (Thermo Fisher Scientific), an Aquilos II cryoFIB/SEM (Thermo Fischer Scientific) or a Crossbeam 550 (Zeiss GmbH). Milling was performed as previously described (Santos *et al*., 2024; Wagner *et al*, 2020). Briefly, inorganic platinum was sputtered on the sample in a Quorum chamber (Quorum Technologies), for the Scios and the Crossbeam 550, or in the main chamber for the Aquilos II. Following this, the cells were covered with a layer of organometallic platinum (Trimethyl [(1,2,3,4,5-ETA.)-1 Methyl-2, 4-Cyclopentadien-1-YL] Platinum) using a gas injection system (GIS). A milling angle of 7-8° relative to the grid’s plane was used for milling, using a stepwise staircase pattern (1nA, 0.5nA, 0.3nA, 0.1nA and 30pA).

### Cryo-ET data collection

Cryo-ET data were collected as previously described (Santos *et al*., 2024). Briefly, ∼230 tilt-series of the sperm neck region, which contains the basal plate, were collected on lamellae using Serial EM PACE-tomo scripts (Eisenstein *et al*., 2023). Human sperm cell data were acquired with a pixel size of 1.63Å using a 300kV Titan Krios G3i Transmission Electron Microscope (Thermo Fisher Scientific) equipped with a Bioquantum energy filter (Gatan) and a K3 direct-electron detector (Gatan). A dose-symmetric tilt scheme was used with increments of 3°, a total dose of 100e/Å^2^ with each tilt 3 e/Å^2^ split in a movie of 6-12 frames, and defocus -2.5 to -5µm.

Cryo-ET data for human fibroblasts was acquired at 1.96Å pixel size using a 300kV Titan Krios G4 (Thermo Fisher Scientific) equipped with a Selectris X energy filter (Thermo Fisher Scientific) and a Falcon 4i direct-electron detector (Thermo Fisher Scientific).

### Tomographic reconstruction, analysis and sub-tomogram averaging

Tilt series were pre-processed using Warp 1.0.9 (Tegunov & Cramer, 2019), followed by tilt series alignment with Aretomo (Zheng *et al*, 2022). Tomograms were reconstructed at ∼10 Å/pix by weighted back projection and lowpass filtered to 60 Å prior to particle picking. Particle picking was carried out with Dynamo (Burt *et al*., 2021; Castano-Diez *et al*, 2012); modelling the sperm basal plate as a surface and sampling it with particle coordinates. Initial model generation and pose refinements were carried out in Dynamo, after which particle coordinates were manually curated in Napari (in-house script). Further pose refinement was then carried out in Relion 3.1 before pose/image, warp/stage, angle refinements in M (Tegunov *et al*, 2021), yielding a 9.4 Å map of the native SUN5-Nesprin3 lattice embedded in the sperm basal plate. For morphometrics analysis, the inner and outer nuclear membranes were segmented either manually in IMOD (J. R. Kremer, 1996) for INM-ONM distance measurements or using MemBrain (Lamm, 2024) for curvedness measurements. A surface morphometrics pipeline (Barad *et al*., 2023) was then used to generate surface meshes of the inner and outer NEs and subsequently measure their curvedness and the distances between these meshes, giving the INM-ONM distance.

### Model building and molecular dynamics flexible fitting (MDFF) of SUN5-Nesprin3 complex

AlphaFold3 (Abramson *et al*., 2024) models of trimeric SUN5 (177-364)-Nesprin3 (34-58) complexes were individually rigid-body fitted into one of the half-maps of the sub-tomogram averaged hexamer using ChimeraX (Pettersen *et al*, 2021) to obtain a model of the SUN5-Nesprin3 hexamer. For flexible fitting, the half-map was zoned around the initial model using phenix.map box (Liebschner *et al*, 2019) with a mask radius of 3 Å. The model of the hexamer was then flexibly fitted into the zoned half-map using NAMDINATOR (Kidmose *et al*, 2019) (200.000 simulation steps, G=0.5, no RSR), improving the model-map cross-correlation from 0.625 to 0.74.

### Cryo-SEM tomography

For volume imaging, cryo-FIB/SEM was performed using a Crossbeam 550 (Zeiss GmbH) (Schertel *et al*, 2013), equipped with a PP3010Z cryo stage and loading system (Quorum Technologies). The exposed surface was milled at an angle of -4° relative to the grid with a current of 700 pA (30 kV) and a slice thickness of 16nm. The pixel size for SEM imaging using an InLens detector at 2.3kV and 50pA was 4.7nm, with a line averaging and scan speed=6. Volumes were processed in Fiji (Schindelin *et al*, 2012). First, contrast-limited adaptive histogram equalisation was used to enhance contrast (Zuiderveld, 1994), then images in the volume stack were aligned using the SIFT algorithm (Lowe, 2004). Horizontal curtain stripes were then removed by horizontal stripe suppression during bandpass filtering. Line segmentation of the *implantation fossa* was carried out manually using Imod (J. R. Kremer, 1996) on six sperm cells.

### Molecular dynamics simulations

The simulations of the structures were performed in two different bilayers. In most cases, for both INM and ONM, we used a heterogeneous bilayer comprising POPC (30%), POPE (30%), POPS (20%), and Cholesterol (20%). In contrast, for the three largest systems (hexamer, trimer of hexamers, and heptamer of hexamers), we used a homogenous membrane (100% POPC). All the systems were constructed using modified scripts from the CHARMM-GUI web server (Wu *et al*, 2014). Sodium and chloride ions were added to achieve an ion concentration of 150 mM. The all-atom CHARMM36m force field was employed for proteins, lipids, and ions, while the TIP3P model was used for water molecules (Best *et al*, 2012; Jorgensen *et al*, 1983; Klauda *et al*, 2010). MD trajectories were analyzed using Visual Molecular Dynamics (VMD), ChimeraX, MDAnalysis, and in-house scripts (Humphrey, 1996; Michaud-Agrawal *et al*, 2011; Pettersen *et al*., 2021).

We performed all simulations using GROMACS 2024 (Abraham *et al*, 2015). The initial setups were energy-minimized for 5,000 steepest descent steps and then equilibrated for 1.5ns in a canonical (NVT) ensemble, followed by seven ns in an isothermal-isobaric (NPT) ensemble under periodic boundary conditions. The restraints on the positions of non-hydrogen protein atoms, initially set at 4,000kJ·mol^-1^·nm², were gradually released during equilibration. The cutoff distance for non-bonded interactions (van der Waals and Coulomb interactions) was set to 1.2nm. Particle-mesh Ewald summation (Darden *et al*, 1993) was employed to handle long-range electrostatic interactions, utilising cubic interpolation and a grid spacing of 0.12nm. The time step was initially 1fs during the NVT equilibration and increased to 2fs during the NPT equilibration. The LINCS algorithm was used to fix all bond lengths (Hess *et al*, 1997). During the equilibration phase, constant temperature and pressure were established using a Berendsen thermostat, combined with a coupling constant of 1.0ps and a semi-isotropic Berendsen barostat with a compressibility of 4.5 ×10^-5^bar^-1^, respectively (Berendsen *et al*, 1984). The Berendsen thermostat and barostat were replaced by a v-rescale thermostat (G. Bussi, 2007) and a C-rescale barostat during the production runs (Bernetti & Bussi, 2020). The list of simulation runs is summarized in Supplementary Table S2.

### Theoretical framework for membrane curvature analysis

The shape of the membrane accommodating the SUN5-Nesprin complex is treated as the Monge surface 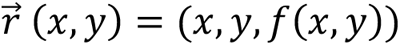, defined by the height function *f*(*x*, *y*). The height function captures the local membrane undulations in three-dimensional space and is parameterised based on x-y coordinates of membrane atoms (Lipowsky, 1991; Mazharimousavi, 2017).

The characterization of topological properties of the membrane surface is feasible through constructing the shape operator, which is the differential of the unit normal vector field 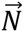 projected on the tangent plane at a surface point. The Mean (*H*(*x*, *y*)) and Gaussian curvatures (*K_G_*(*x*, *y*)) of the surface can be derived from the invariants of the shape operator matrix, which depends only on the height function (Steigmann, 2017).

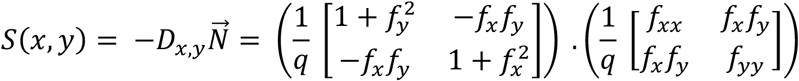

Where 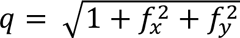 and *f_x_* and *f_xx_* denote 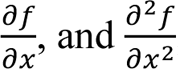, respectively; similar definition applies for the y-direction. Consequently, the determinant and half of the trace of the *S*(*x*, *y*) matrix corresponds to the Mean (*H*(*x*, *y*)) and Gaussian curvatures (*K_G_*(*x*, *y*)) of the membrane (Steigmann, 2017).

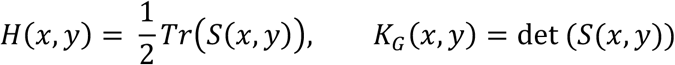

To achieve this, we approximated the height function of the membrane surface, parameterizing through simulation trajectory data using a Python script adapted from Bhaskara et al. (Bhaskara *et al*., 2019) . In their approach, the height function was expressed as a two-dimensional Fourier expansion, and they minimized the squared deviations between the Fourier-constructed membrane heights and the observed values. To ensure the consolidation and reproducibility of our data, each molecular dynamics simulation is repeated three times using replicated setups, each starting from randomized initial velocities. Curvature measurements, whether contour maps or the time evolution of local mean curvature, are averaged across all replicas (Abramson *et al*., 2024; Pettersen *et al*., 2021)(Kidmose *et al*., 2019).

### Estimation of expected curvature of SUN5-containing flattened area

To understand the effect of SUN5 lattice on the membrane remodelling, we try to estimate the initial expected curvature of the SUN5-containing region in the absence of the SUN5 lattice. For this, we developed a computational method using Python (v3.12) with the NumPy and SciPy libraries. The process began with coordinate data derived from manual tracing of membrane profiles on the SEM images using image analysis software (IMOD) (J. R. Kremer, 1996). Three distinct profiles were traced: the flattened membrane of interest, and the adjacent membranes to its left and right.

From the full set of traced points, three points (start, middle, and end) were selected to serve as anchor points. These anchors were used to define two idealised curves: 1) an ‘ideal expected curve’ representing the anticipated path of the membrane without flattening, and 2) an ‘ideal flattened curve’ representing a perfectly linear version of the flattened segment. A parametric B-spline was independently fitted to each set of anchor points using the splprep function from SciPy’s interpolation module. This approach provides a smooth, continuous representation for both the expected and flattened ideal membrane shapes.

Flattening was quantified using two metrics. First, to derive a direct measure of local curvature, we calculated the curvature (κ) for the splines using the standard formula from differential geometry:

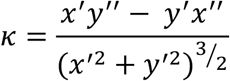

where x′, y′ and x′′, y′′ are the first and second parametric derivatives of the spline, respectively. The average curvature was calculated for the segment of the ‘ideal expected spline’ corresponding to the flattened region, as well as for the ‘ideal flattened spline’.

Second, the deviation of the actual traced membrane from the ideal state was measured by calculating the maximum perpendicular distance (h) of the fitted ‘ideal expected spline’ to the traced flattened profile. Then, using the length of the flattening region (L), we estimated the radius of circle fitting into the ideal expected line using:

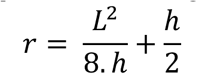

Then local curvature was calculated as:

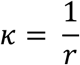

### Immunostaining

For immunostaining of human skin fibroblasts cells were seeded on #1.5 glass coverslips for 24h at 37°C, 5% CO_2_ before fixation in 4% PFA-PBS for 10 minutes at room-temperature (RT). Cells were then washed in PBS and quenched with 50mM NH_4_Cl-PBS for 15 minutes at RT, followed by washes in PBS. Cells were then simultaneously permeabilised and blocked in 3% bovine serum albumin (BSA), 0.2% Triton-X100 in PBS for 30 minutes at RT. Following this, cells were incubated for 2h at RT with primary antibodies (rabbit anti-SPG4L, Invitrogen PA5-61761; mouse anti-Nesprin3, Proteintech 68420-1-IG) diluted in 3% BSA, 0.2% Triton X-100 in PBS. Cells were washed in0.5% BSA, 0.05% Triton X-100 in PBS and incubated for 1 hour at RT with fluorescently labelled secondary antibodies (donkey anti-rabbit Alexa Fluor 647, Abcam ab181347; donkey anti-mouse Alexa Fluor. 488, Abcam ab181289) diluted in 3% BSA, 0.2% Triton X-100 in PBS. Cells were then washed in 0.5% BSA, 0.05% Triton X-100 in PBS, followed by PBS and mounted using Prolong Gold (Thermo Fischer Scientific).

### Structured Illumination Microscopy

SIM imaging was performed using a Zeiss Elyra S.1 system (Carl Zeiss) using a 63x 1.46NA plan-apochromat oil immersion objective (Carl Zeiss), with 405nm, 488nm, 561nm and 647nm laser lines and a CMOS camera. Z-stacks were collected every 91nm through the height of the nucleus, with 5 rotations and frame averaging of 4. The affine method was used for channel alignment calibration, using mounted TetraSpeck beads (Invitrogen). Data analysis, including channel alignment, deconvolution and SR-SIM post-processing, was performed using Zeiss Zen Black 2.3 software (Carl Zeiss).

### Immunohistochemistry

Glass slides with pre-mounted FFPE-embedded testis sections (5μm thick) were deparaffinised and rehydrated before heat-induced antigen retrieval in 10mM Sodium Citrate, 0.05% Tween 20 (pH 6). Following antigen retrieval, tissue sections were washed in 0.05% Triton-X100 in TBS for 15min with rocking at room temperature, followed by incubation in blocking solution (1%BSA, 10% Donkey Serum in TBS) for 30min at room temperature. Tissue sections were then incubated with primary antibody rabbit anti-SPAG4L (Invitrogen, PA561761) diluted in 1% BSA in TBS, overnight at 4°C. Tissue sections were then washed in 0.05% Triton X-100 in TBS, for 10min at room temperature, with rocking, 3 times and incubated for 1h at room temperature with fluorescently labelled secondary antibody (donkey anti-rabbit Alexa Fluor 647, Abcam ab181347; or donkey anti-rabbit Alexa Fluor 488, Abcam 181346), diluted in 1% BSA in TBS. Tissue sections were washed in 0.05% Triton-X100 in TBS for 10min at room temperature three times, followed by one TBS wash and one final wash in diH_2_O. Tissue sections were stained with Hoechst 33342 (Thermo Fischer Scientific) and mounted using #1.5 coverslips and Prolong Gold Mounting media (Thermo Fischer Scientific). (Santos *et al*., 2024).

### Confocal Microscopy

Testis sections were imaged using an Andor Bench Top Confocal BC43 (Oxford Instruments), equipped with a 60x 1.42NA oil immersion objective, 405nm, 488nm, 561nm and 638nm lasers and an sCMOS detector. Imaging acquisition was performed using Fusion software (Oxford Instruments), and image analysis was performed using Imaris software (Oxford Instruments).

---

**Fig S1.**
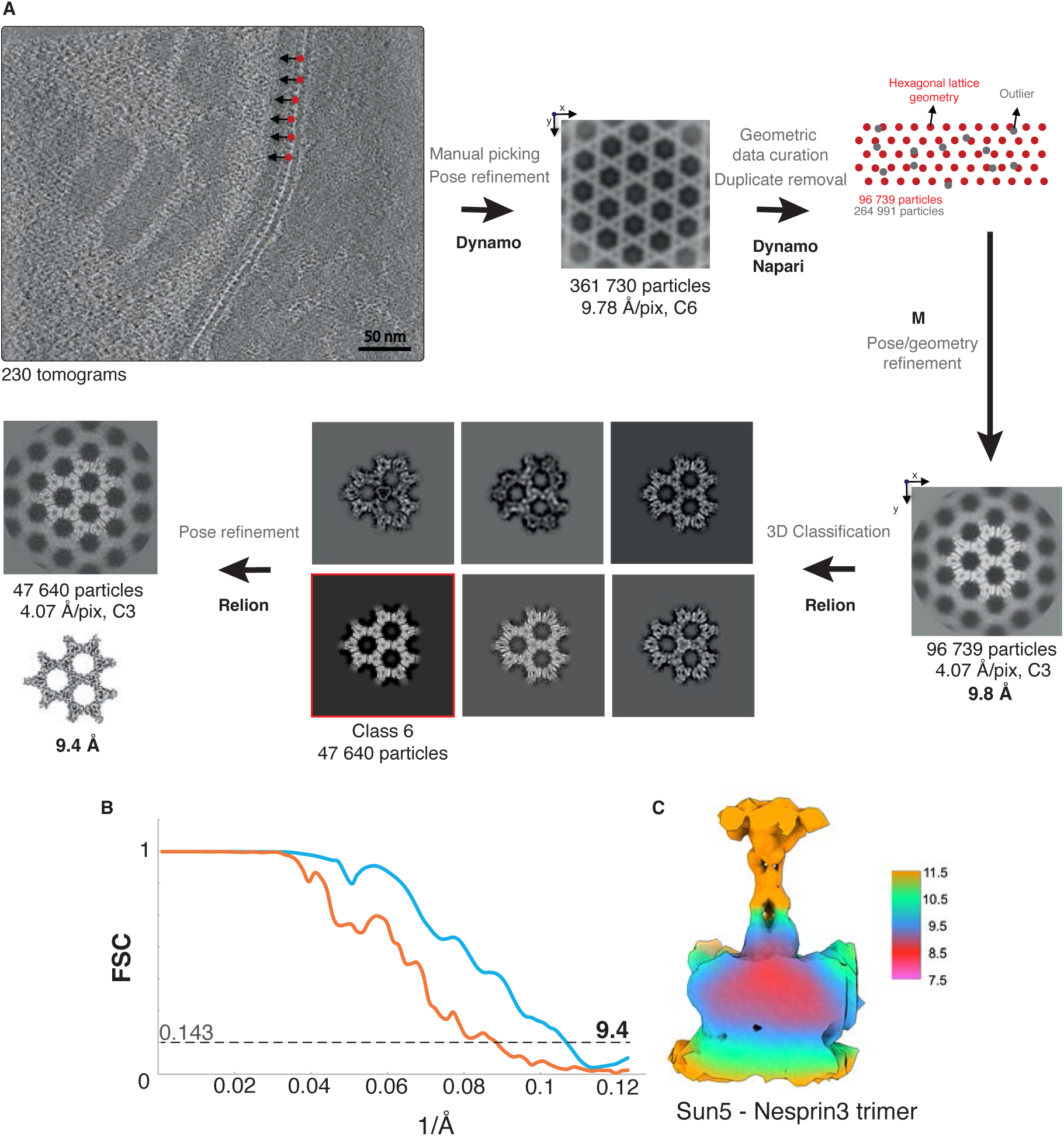
Fourier Cell Correlation and local resolution. **A**) Schematic of the sub-tomogram averaging pipeline. Particles were picked in Dynamo, on tomograms reconstructed with Warp at 9.78 Å/pix. Sub-tomogram volumes were aligned with Dynamo, and particles positions were curated manually in Napari using an in-house script, based on lattice geometry. Poses were refined in M, with C3 symmetry imposed, as well as tilt series geometry, resulting in a sub-tomogram average of the hexagonal SUN5-Nesprin3 lattice at 9.8 Å. Iterative 3D classification and subsequent 3D refinement in Relion 3.1 resulted in a 9.4 Å average from 47,640 particles. **B)** Fourier Shell Correlation of the final SUN5 trimer sub-tomogram average, showing the masked (blue) and unmasked (orange) curves. **C**) Final SUN5-Nesprin3 trimer sub-tomogram average, colour coded according to local resolution.

**Fig S2.**
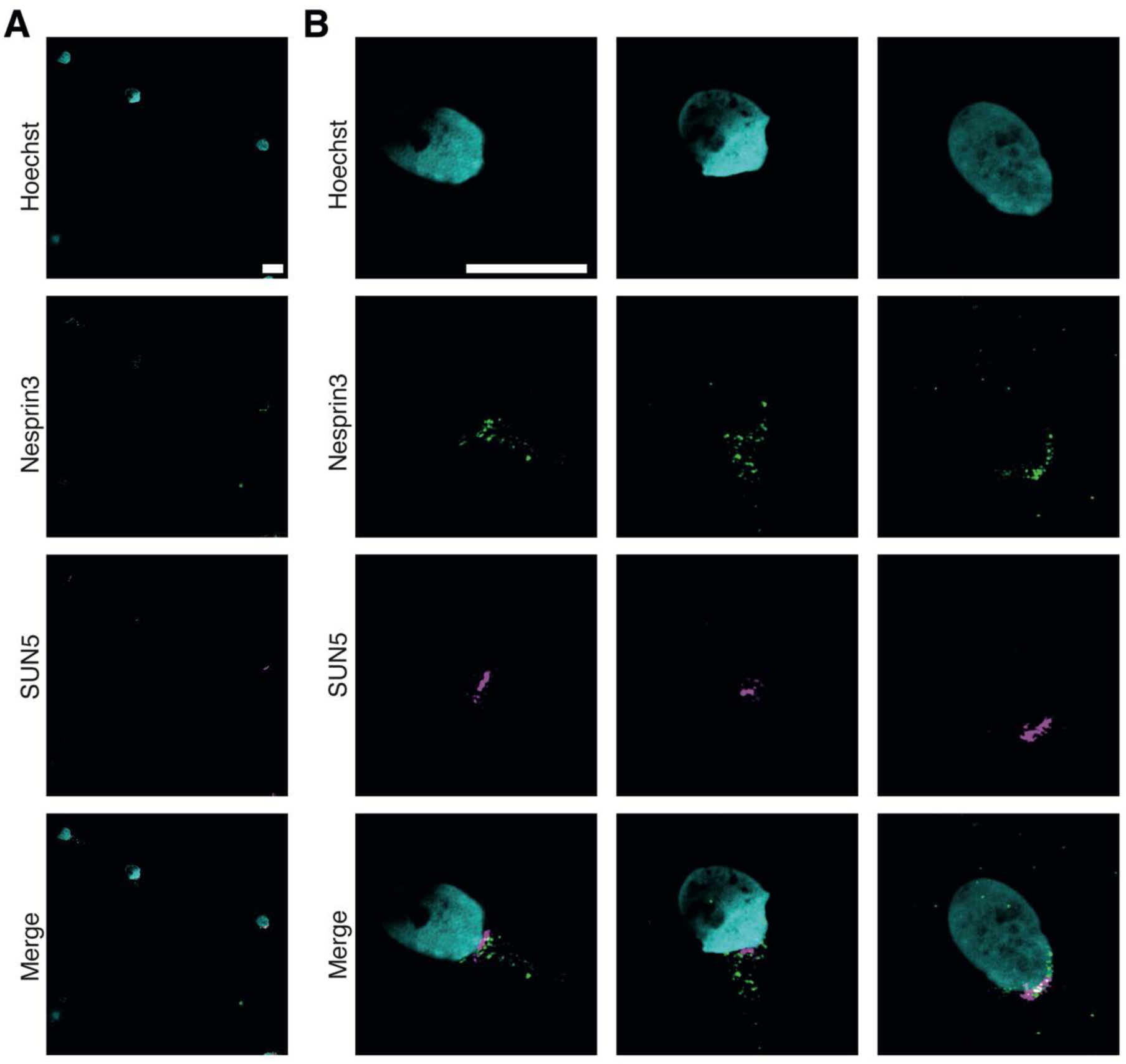
SUN5 and Nesprin3 localisation in mature human sperm cells. **A**) Structured Illumination Microscopy image (SIM) showing an overview of mature human sperm cells. Immunofluorescent labelling for SUN5 is shown in magenta, for Nesprin3 in green, and DNA stain Hoechst is shown in cyan. Scale bar=5um. **B**) Detail of three human sperm cells from the image shown in A). Scale bar=5um.

**Fig. S3.**
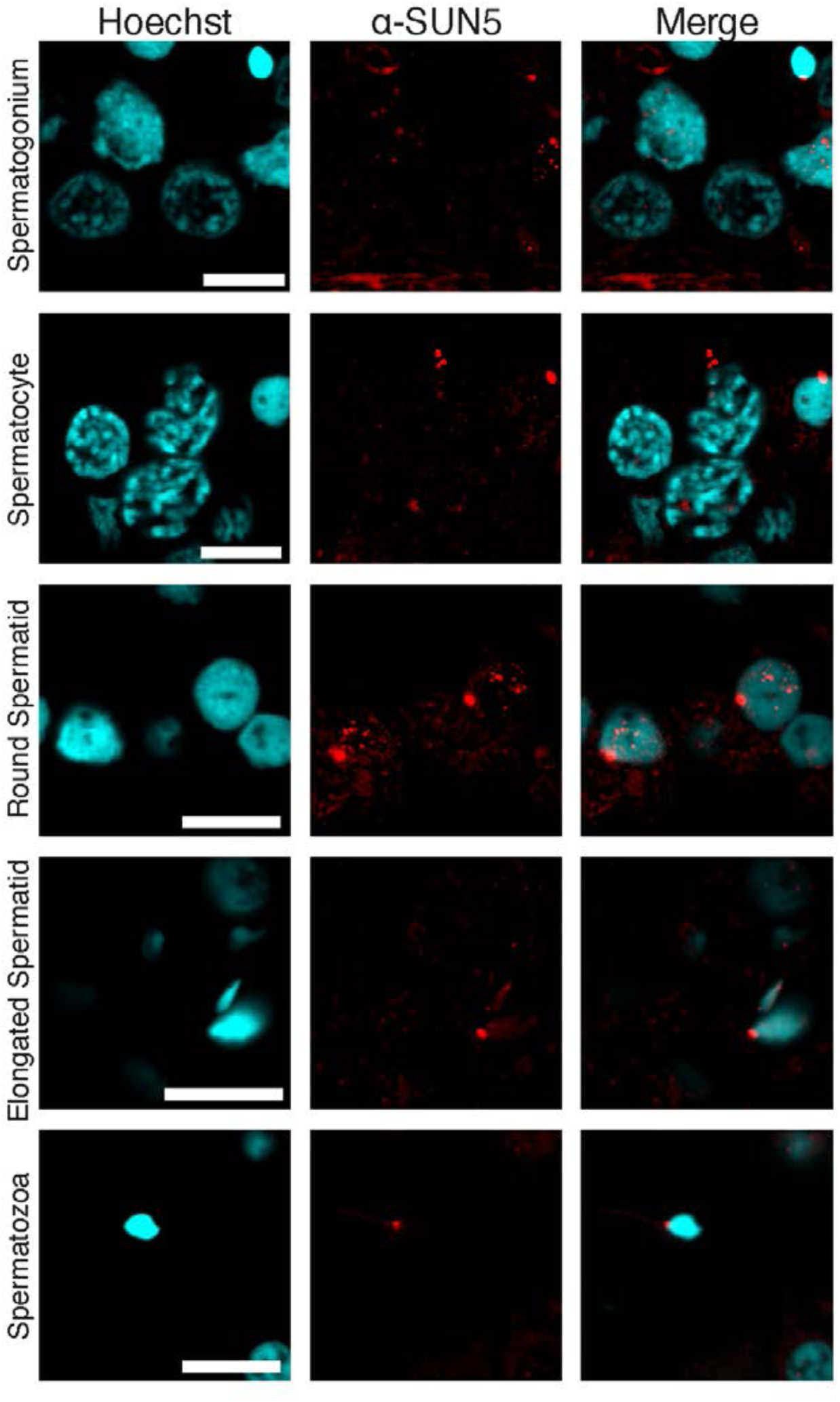
SUN5 expression and distribution of healthy human testis tissues. **A**) Immunohistochemical staining against SUN5 is shown at the different stages of sperm differentiation, from spermatogonia to spermatozoa. SUN5 is shown in red, and DNA stain Hoechst is shown in cyan. Scale bar=10um.

**Fig. S4.**
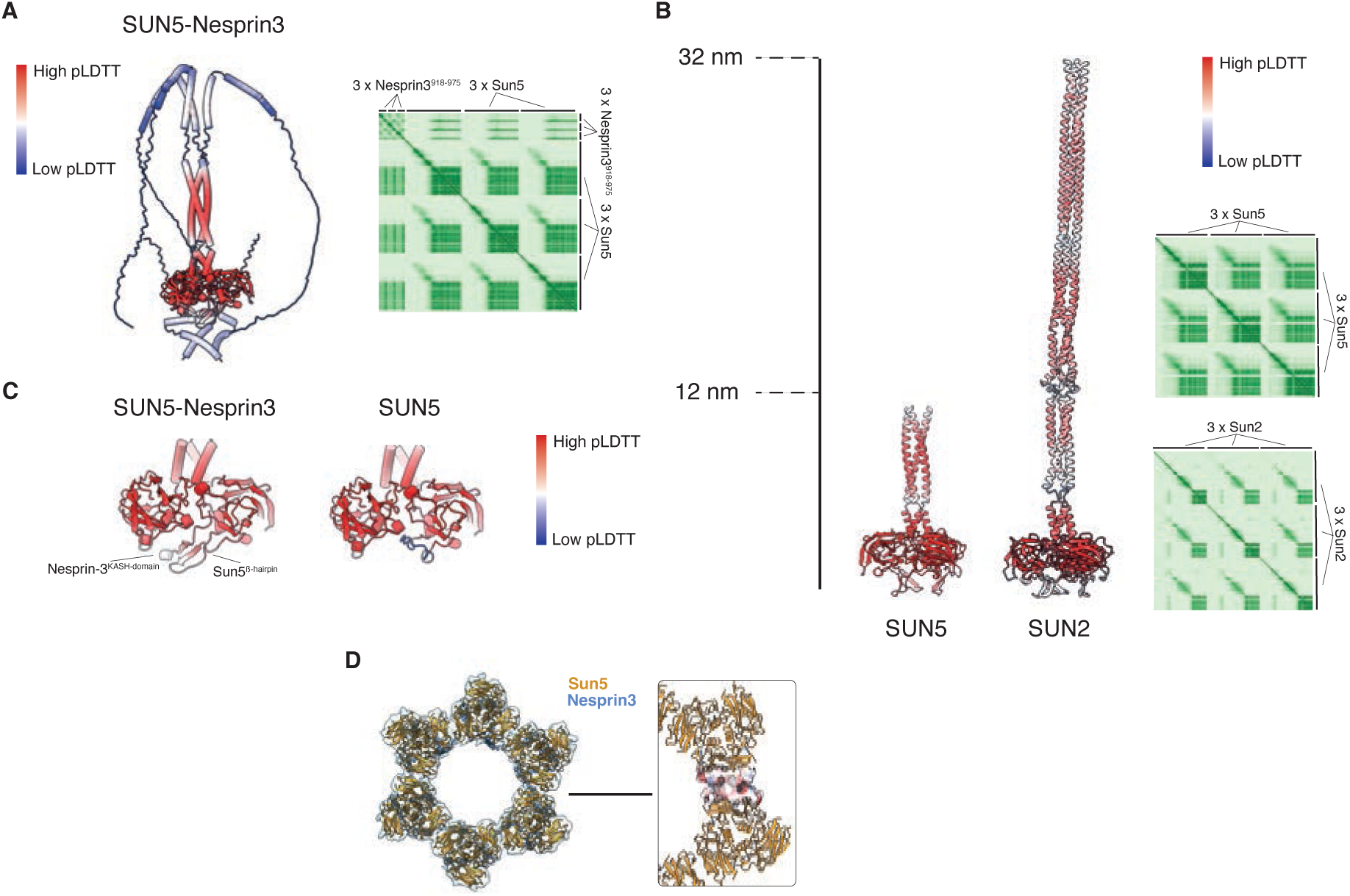
Alpha-fold predictions and inter-trimer interface. **A**) AlphaFold prediction of a SUN5-Nesprin3 trimer, coloured according to pLDTT score, including PAE plot. **B**) Comparison of trimeric SUN5 and SUN2 predictions, coloured according to the pLDTT score, showing a 12nm long triple coiled coil domain for SUN5 and a 32nm long coiled coil domain for SUN2. **C)** Comparison of a trimeric SUN5-Nesprin3 prediction and a trimeric SUN5 prediction, coloured according to the pLDTT score, revealing an exposed and folded SUN5 KASH-lid only when Nesprin3 is bound. **D**) Cross-section view of a hexa-trimer at the level of the SUN domain with the fitted model. On the right zoomed-in inset showing the polar lattice inter-SUN5 trimer interface as an electrostatic surface.

**Fig S5.**
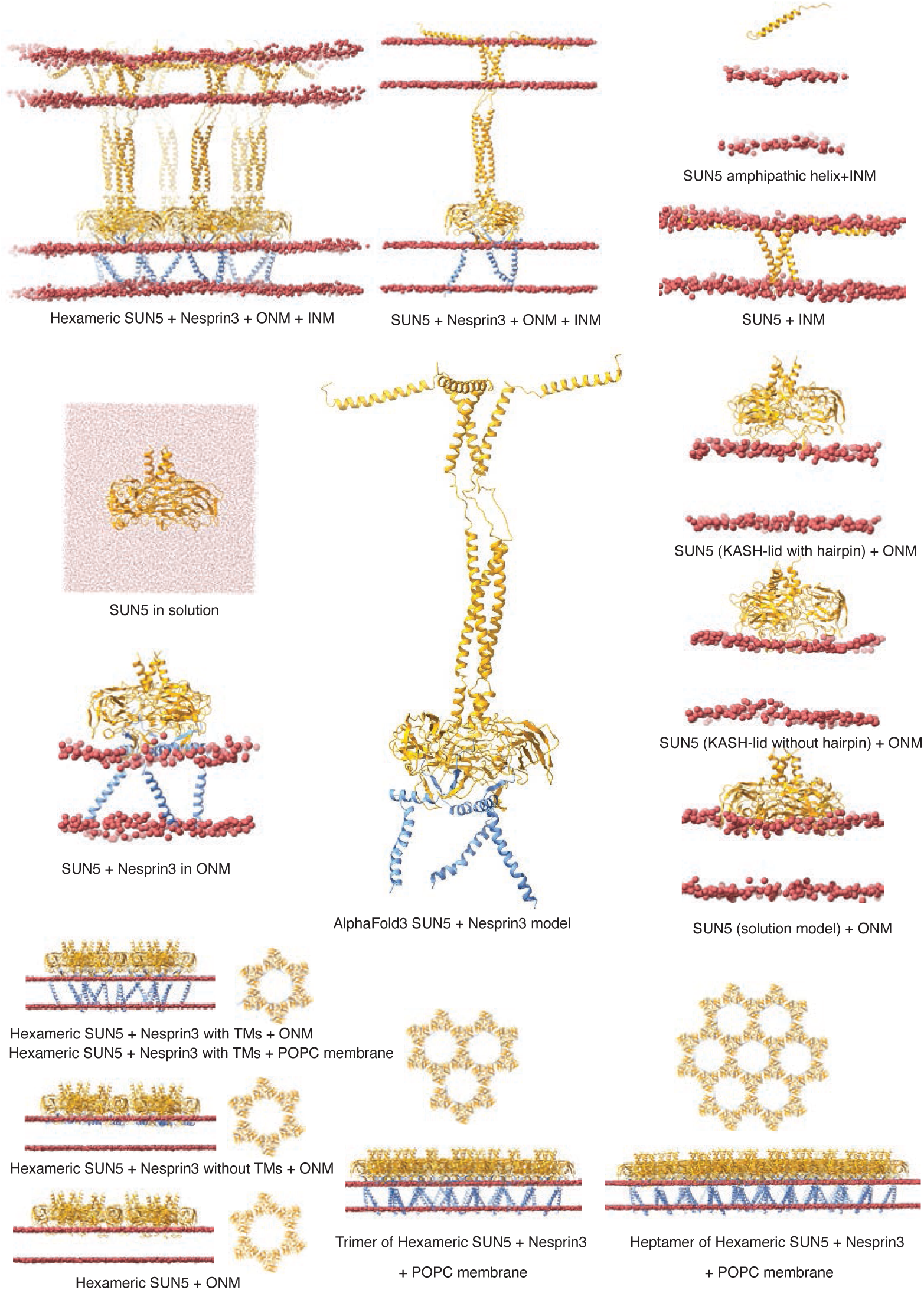
The schematic figure represents all the simulations performed in this study. SUN5, Nesprin3, and Membrane (P atoms) are colored orange, blue, and red, respectively. ONM, INM, and TM stand for outer nuclear membrane, inner nuclear membrane, and transmembrane region, respectively. In all simulations, INM and ONM have a similar composition (30% POPC, 30% POPE, 20% POPS, and 20% Cholesterol) except for three simulations that were performed in a homogenous POPC membrane (hexamer, trimer of hexamers, and heptamer of hexamers).

**Fig. S6.**
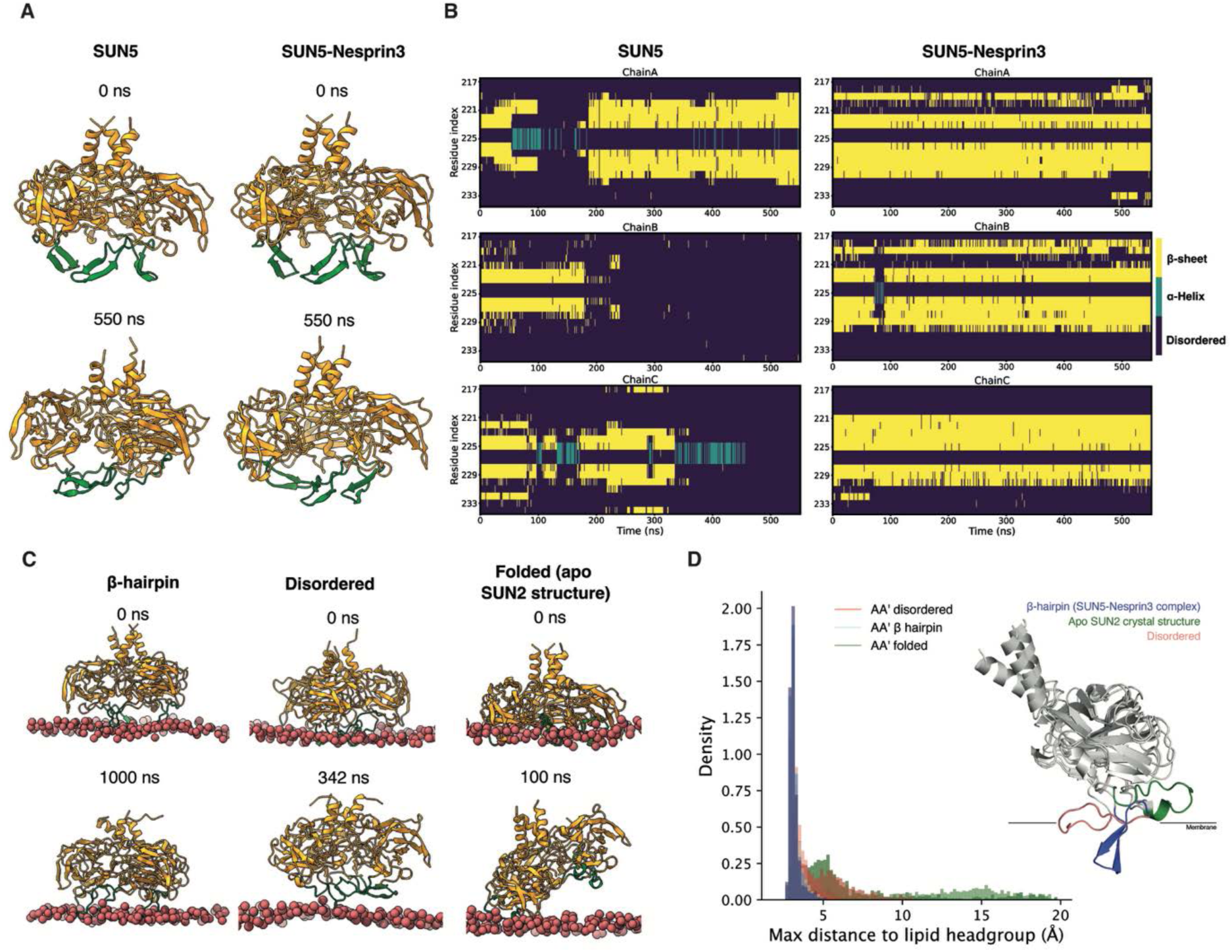
MD simulation of the SUN5-domain interactions with the SUN5 KASH-lid in different secondary structure conformations. **A**) MD simulation of SUN5 alone, and with Nesprin3 in solution. **B**) Changes in secondary conformation of KASH-lid during the simulation of SUN5 alone and with Nesprin3 in solution. **C**) MD simulation of the SUN5-domain interaction with the ONM with the SUN5 KASH-lid in different secondary structures (β-hairpin (left), disorder (middle), and folded (right)). **D**) The distribution of the distance of SUN5 from the membrane during the simulations of the SUN5-domain interaction with the ONM, with the SUN5 KASH-lid in different secondary structures.

**Fig. S7.**
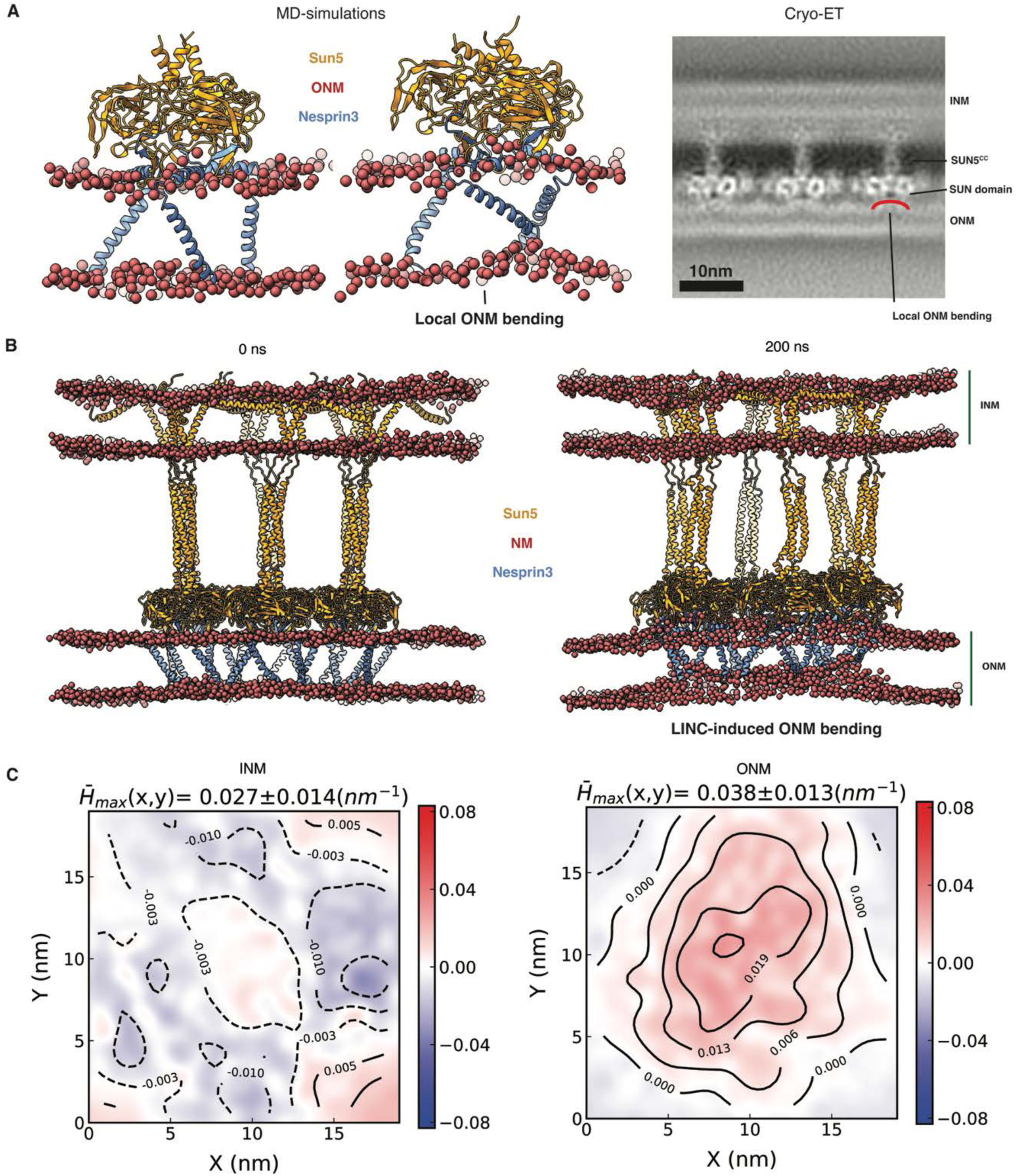
The *curvature-inducing* property of the SUN5-Nesprin3 complexes. **A**) Atomistic MD simulations of the trimeric SUN5-Nesprin3 perturb the membrane locally (left), similarly to what is observed in the cryo tomographic data (right, INM stands for inner nuclear membrane and ONM for outer nuclear membrane). **B**) The full-length SUN5-Nesprin3 complex in the NE can induce curvature in the ONM during the MD simulation timescale. **C**) Contour maps of the averaged curvature profiles for the ONM (right) and INM (left) show the extent of curvature induction around the centre of the SUN5-Nesprin3 hexamer complex.

**Fig. S8.**
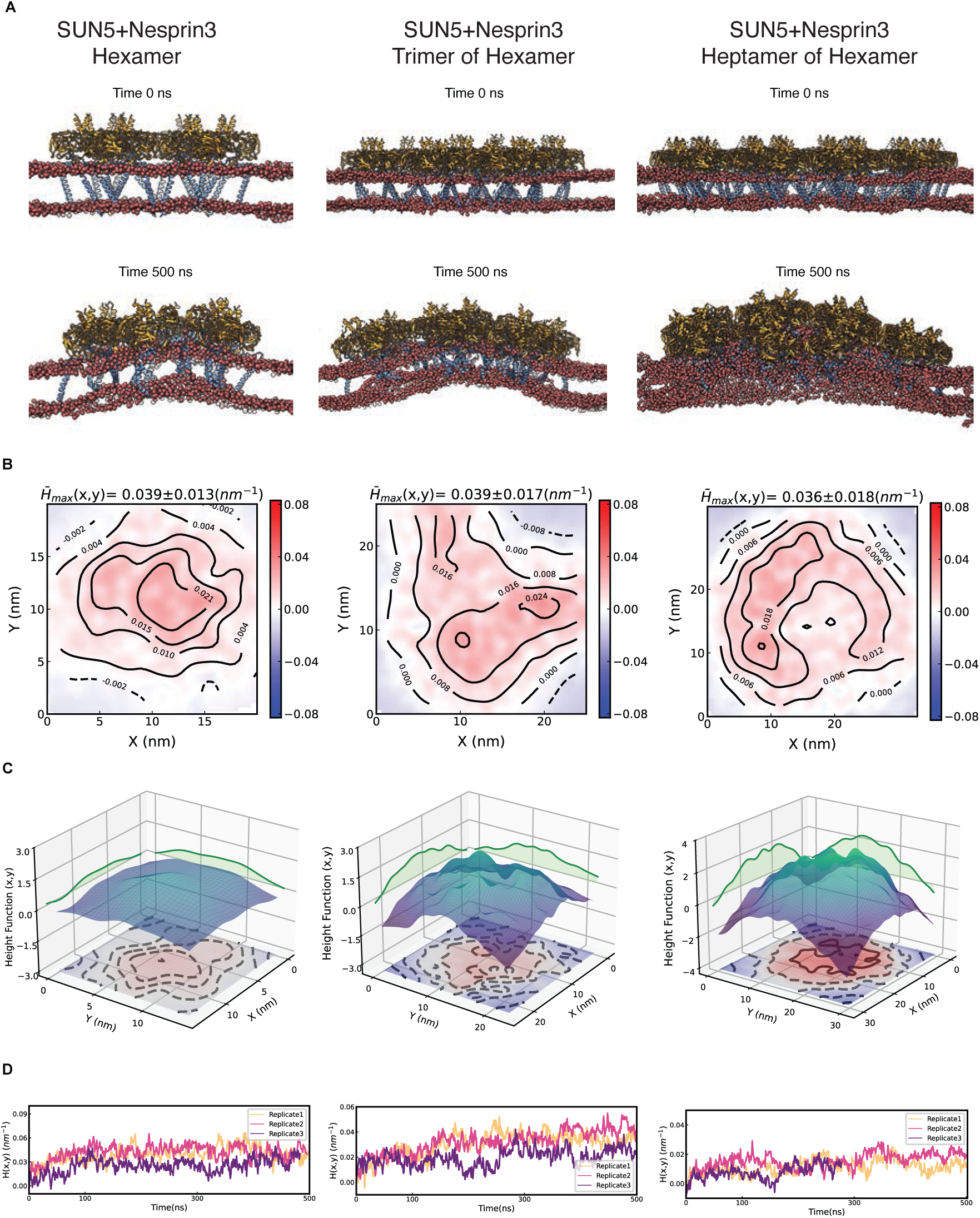
The *curvature-inducing* property of the SUN5-Nesprin3 complexes in different oligomeric states. **A**) SUN5-Nesprin3 hexamer (left), trimer of hexamers (middle), and heptamer of hexamers (right) induce global curvature in the bilayer structure. Side view snapshots at the beginning of the simulations (top panels) and the end of the simulations (bottom panels) with SUN5, Nesprin3, and Membrane (P atoms) coloured orange, blue, and red, respectively. **B**) Contour maps of the averaged curvature profiles for each system show the extent of curvature induction around the center of the SUN5-Nesprin3 complex. The maximum value of curvature fields and Std. is shown at the Top. **C**) Local membrane shape is approximated by a height function of the midplane (height contour map along the plane). **D**) Time-resolved averaged curvature profiles for the replicates of each system.

**Fig. S9.**
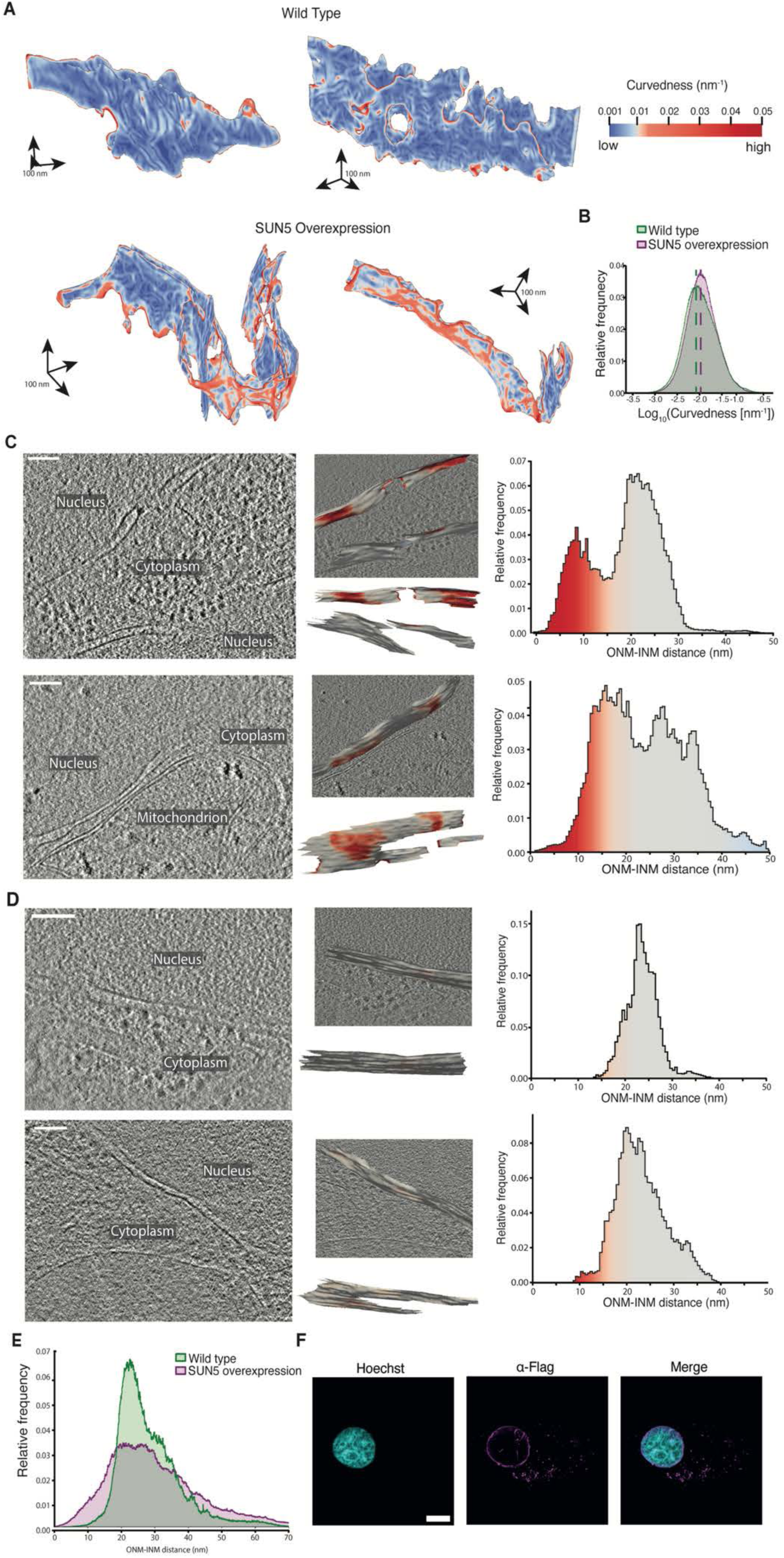
Morphometric analysis of the NE in human fibroblasts overexpressing SUN5. **A**) Example tomographic segmentations of the ONM and INM for analysis of membrane curvedness, coloured using a heatmap of membrane curvedness (top right). The top panel shows a typical wild type segmentation, and the lower panel shows a typical (65% of the tomograms) SUN5 overexpression segmentation. **B**) Combined histogram showing nuclear membrane curvedness across all the analysed tomograms of wild-type (green) and SUN5 overexpressing (purple) cells, plotted on a log scale. Dashed vertical lines show the peak histogram values. A total NE area of 8.27 μm^2^ over 17 tomograms from 11 cells was analysed for the wild type, and 5.82 μm^2^ over 15 tomograms from 9 cells was analysed for the SUN5 overexpression. **C**) 2D slices of reconstructed tomograms of SUN5 overexpressing cells (left). Segmentations of the ONM and INM for morphometrics analysis are shown in the central panels, coloured using a heatmap of the ONM-INM distance. The distribution of ONM-INM distance for each tomogram is plotted (right), with the histograms overlaid with the same heatmap. The distributions show two peaks representing nuclear envelope patches at ∼10nm and ∼25nm ONM-INM distance. **D**) As in C) but from wild-type cells. The distributions show a single peak at ∼20 nm ONM-INM distance. Scale bar=100nm. **E**) Combined histogram showing the distribution of ONM-INM distances across all the analysed tomograms. To compare the ONM-INM distribution between wild-type (green) and SUN5 overexpressing (purple) cells, a Levene’s test was used after calculating the variance for each tomogram. This showed that the ONM-INM distance is significantly more variable in SUN5 overexpressing cells compared to the control cells (Levene test statistic = 6.543, p = 0.017). **F**) Confocal image of human fibroblasts transfected with Flag-SUN5 immunolabelled with anti-FLAG antibody (magenta). DNA is shown in cyan (Hoechst). Scale bar=10μm.

**Fig. S10.**
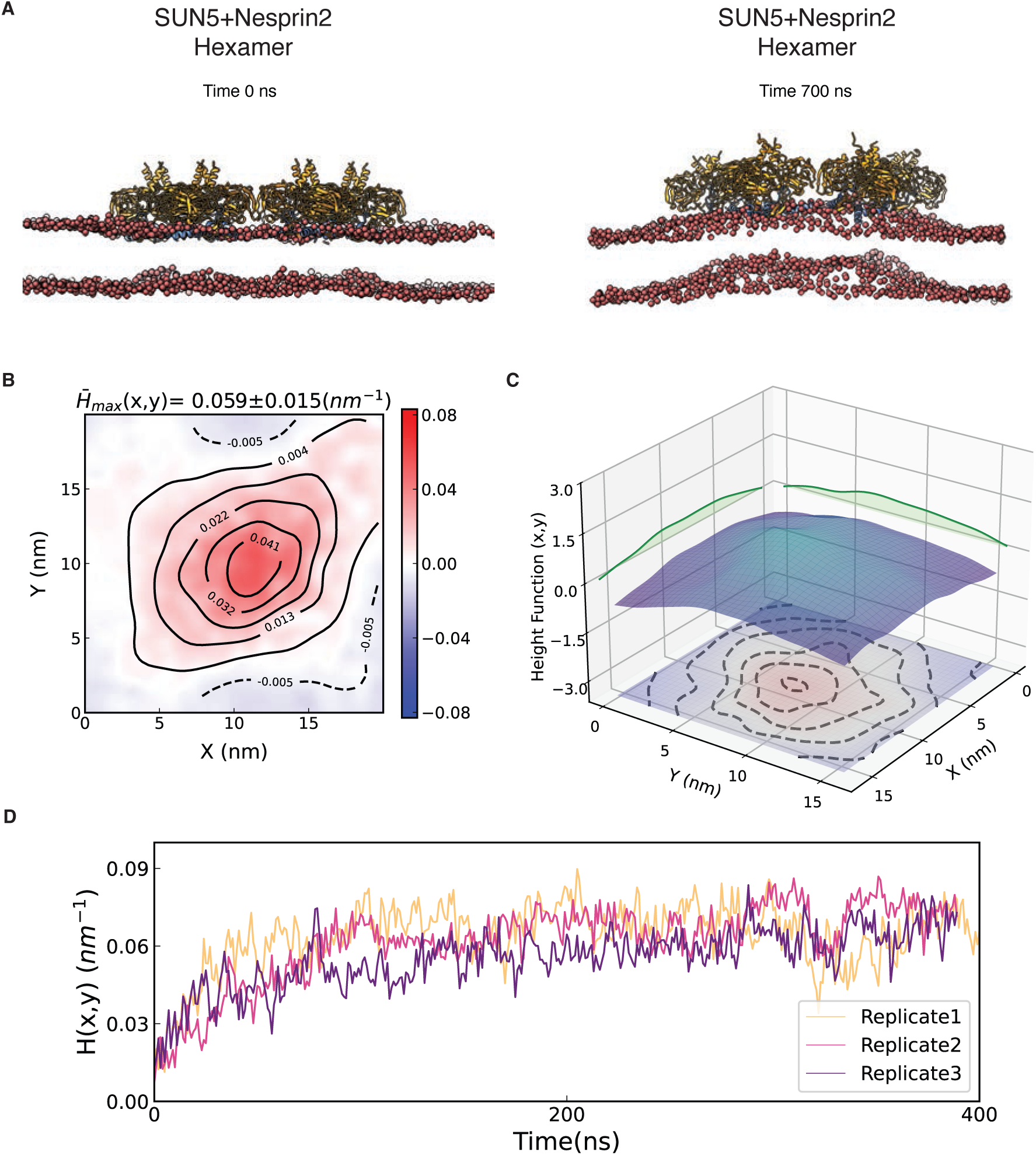
The *curvature-inducing* property of the SUN5-Nesprin2. **A**) SUN5-Nesprin2 hexamer induces global curvature in the bilayer structure. Side view snapshots at the beginning of the simulations (top panels) and the end of the simulations (bottom panels) with SUN5, Nesprin2, and Membrane (P atoms) coloured orange, blue, and red, respectively. **B**) Contour maps of the averaged curvature profiles for each system show the extent of curvature induction around the center of the SUN5-Nesprin2 complex. The maximum value of curvature fields and Std. is shown at the Top. **C**) Local membrane shape is approximated by a height function of the midplane (height contour map along the plane). **D**) Time-resolved averaged curvature profiles for the three replicates.

**Table S1.**

**Result of the expected curvature of the SUN5 region obtained from estimating curvature flattening in the SEM images.** Method 1: calculating the curvature (κ) for the splines. Method 2: calculating the curvature using the maximum perpendicular distance (h) of the fitted ‘ideal expected spline’ to the traced flattened profile.

**Table S2.**

**Details of the atomistic Molecular Dynamics simulation setups performed in this work.**

## Supplementary text

### SUN5 amphipathic helix

Although AlphaFold did not confidently predict the structure of the SUN5 amphipathic helix (aa 65-85), the following lines of evidence support its existence and association with the membrane. Analysis using the Positioning of Proteins in Membrane (PPM 3.0) program indicates the helix orients parallel to the membrane and sits on top of the lipid head groups (Lomize *et al*, 2022). This localization is corroborated by molecular dynamics simulations, which show the helix interacting with the membrane and integrating with the membrane’s head group region within 150ns.

### Role of SUN5 KASH-lid and Nesprin3 in the ONM anchoring of the SUN domain

We performed atomistic MD simulations of the SUN domain in both solution and an ONM lipid bilayer, with and without the Nesprin3 KASH domain (Table S2).

In simulations of the SUN domain in solution, the KASH-lid β-hairpin was stable only in the presence of Nesprin3. In its absence, the KASH-lid became unstable and mostly folded back into the SUN domain, presumably to shield its hydrophobic residues from the aqueous solvent (Fig. S6A-B).

We performed four distinct simulations with the ONM to test the stability of the membrane-anchored state (Table S2) (Fig. S6C-D).

1. In the presence of the Nesprin3 KASH domain, the SUN domain remained stably bound to the ONM.
2. When simulating the SUN domain alone but starting with the KASH-lid already in its pre-formed β-hairpin conformation, it also remained stably bound, and the KASH-lid kept its β-hairpin conformation.
3. In contrast, when the SUN domain was simulated alone with the KASH-lid in a disordered conformation (taken from solution simulations), it partially dissociated from the ONM.
4. Finally, when the KASH-lid was modelled in its fully folded-in state (based on the SUN2 structure, PDB ID 3UNP), the SUN domain completely dissociated from the membrane.

Our results reveal a crucial and nuanced role for Nesprin3 in this process. Consistent with our AlphaFold3 predictions, we found that Nesprin3 binding is essential for achieving a *stable* β-hairpin conformation. In simulations of the SUN domain in solution, the KASH-lid β-hairpin was unstable in the absence of Nesprin3, while in the simulations in the ONM environment, it remained stable. Therefore, the process of anchoring the SUN domain exhibits molecular hysteresis, acting like a one-way switch: Nesprin3 is required for folding and insertion of the KASH-lid into its stable, membrane-bound β-hairpin conformation. However, once this state is achieved and anchored within the stabilising lipid environment of the ONM, it is self-maintaining and no longer requires the presence of Nesprin3 to prevent dissociation. It is as if the KASH-lid “remembers” this state with the help of its membrane environment.

### Effect of membrane and protein complex composition on the curvature-inducing property of the SUN5-Nesprin3 complex

#### The effect of membrane composition

We conducted simulations on two different membranes: a heterogeneous bilayer (PC/PE/PS/CHOL) and a pure POPC one. We primarily used the pure POPC for curvature scaling analysis across various lattice sizes. The main reason for this simplification was to reduce computational resources, as larger systems (trimer and heptamer of hexamers) required longer simulation times for proper equilibration and lipid mixing in a heterogeneous bilayer. Furthermore, employing a membrane with homogeneous lipid composition minimises compositional complexity, ensuring that any observed changes by protein can be pinpointed more easily. However, simulating one of the systems (hexamer cluster of SUN5+Nesprin3) in both membranes allowed us to compare the effect of membrane composition. Our results showed that the SUN5+Nesprin3 hexamer induces higher curvature in a heterogeneous membrane (PC/PE/PS/CHOL) than in a pure PC bilayer. The enhanced curvature can be attributed to a change in the collective mechanical properties of the membrane. Specifically, cholesterol is well-known to modulate membrane fluidity and bending rigidity, and its presence in the heterogeneous system likely makes the membrane more permissive to deformation by the protein lattice.

#### The effect of double membrane

We also simulated the full-length SUN5+Nesprin3 hexamer within a double membrane system, which revealed a reduction in remodelling capability compared to single-membrane simulations. The curvature induced by the protein lattice on the outer nuclear membrane was smaller, and no discernible deformation was observed on the inner membrane within the simulation timescale. This attenuated effect can be attributed to several factors: 1) A coupled double membrane system probably has the substantially higher mechanical load and bending rigidity; 2) The flexible nature of the SUN5’s inter-membrane linker domain results in the inefficient force transduction, which likely dissipates force through conformational changes; and 3) The possibility that the a single hexamer used here was insufficient to generate the cooperative force required to overcome the combined energetic barriers to bend both membranes together.

### The link between the size of the SUN5 oligomer and its induced curvature

Our simulations revealed a non-linear, inverse relationship between the size of the protein oligomer (L) and the magnitude of the principal curvature (k₁) it induces at the center point of the SUN5 oligomer. A log-log plot of this relationship yielded a linear trend, indicating that the induced curvature scales with lattice size (L) as a power law (k_1_ ∝ L^−α^, where α is the positive scaling exponent). This finding suggests that the curvature is probably determined by the collective mechanics of the protein-membrane system, where the adequate bending rigidity increases with the size of the protein lattice.

This observation can be understood by considering the protein lattice and the underlying lipid bilayer not as separate entities, but as a coupled, composite material. The total energy of this system is dominated by the elastic energy required to bend this composite sheet, which scales with its area (A) and the square of its curvature (E_bend_ ∝ A ⋅ k_1_^2^). In this model, the behaviour at different lattice sizes is as follows: *Small Lattices*: A small, localised protein patch can readily impose its preferred intrinsic curvature on the membrane. The energy cost to bend this small area is minimal, and the surrounding lipid bilayer is compliant enough to accommodate the local deformation. *Large Lattices*: As the lattice grows, it forms a large, continuous sheet that is most likely stiffer than the bare membrane and also the smaller oligomers. The effective bending rigidity increases with the lattice area (L). Consequently, the energetic penalty for bending the entire, large patch to a high curvature becomes prohibitively expensive. To minimise the total free energy, the system adopts a state of lower overall curvature. It is energetically more favourable for the large, stiff lattice to remain relatively flat than to pay the immense energy cost required to bend the entire structure. The observed power law, where curvature decreases with lattice size, is the macroscopic signature of this stiffness-dominated mechanism. The relationship is non-linear because the energetic trade-off is not a simple additive process. This demonstrates that the protein assembly’s primary role in this regime is to form a rigid scaffold that, as it grows, progressively resists the high-energy cost of deformation, leading to a flatter overall structure. This finding can explain the lattice’s observed biological function. In the sperm, the SUN5 lattice forms on a pre-existing, inversely curved inner/outer nuclear membrane. The high bending rigidity for the larger lattices predicted by our simulations is exactly the property that allows them to serve as effective flattening agents. A very stiff object will strongly resist deformation into a shape that differs from its low-energy, relatively flat state. Therefore, the higher stiffness of the larger lattices enables them to easily overcome the membrane’s initial negative curvature and impose their own flatter shape, providing a clear mechanism for the experimentally observed membrane remodelling.

It is important to note that these simulations were performed using a membrane patch of finite size, which was not significantly larger than the protein lattices. This computational constraint introduces boundary effects that contribute to the effective stiffness of the system. While these finite-size effects may influence the exact value of the scaling exponent (α), the qualitative trend and the underlying physical conclusion that system rigidity increases with lattice size remain robust. Furthermore, we performed a coarse-grained (MARTINI) simulation of hexameric SUN5-Nesprin3 in a bigger patch (45×45 nm compared to 34×34 nm of atomistic), and we observed very similar curvature (CG data are not shown in this paper).

### Shape of the SUN5 lattice

An important observation is the change in the shape of the SUN5 oligomer and the resulting curvature. For the hexamer, the main curvatures were anisotropic, with k₁ consistently larger than k₂. This reflects the influence of the individual hexamer, which gradually becomes structurally elongated during the simulation and thus has a preferred bending axis. Although the reason for this shape anisotropy is not clear, it was seen in all hexameric simulations regardless of the hexamer composition. However, as the lattice size increases, the principal curvatures converge (k₁ ≈ k₂), suggesting that the lattice becomes mechanically isotropic. This likely happens because the local anisotropies of the individual hexamers are averaged out in the larger, more cohesive structure. In a large lattice, the overall properties overshadow the shapes of individual hexamers and edge effects, leading to a more uniform, symmetrical mechanical response.

### Estimating curvature flattening in the SEM images

We use a computational method to estimate the amount of curvature flattening in the SUN5-containing region of sperm cells (See the Materials and Methods section). While our approach offers a quantitative measure of membrane flattening, it has several inherent limitations. The initial identification of flattened and adjacent membrane profiles depends on manual tracing from cryo-electron tomograms, which can introduce user-dependent variability. Additionally, modelling the “ideal expected curve” with a B-spline is an approximation that probably does not reflect the actual initial state of the membrane. Another key assumption of our “bridging spline” method is that membrane regions adjacent to the SUN5 lattice are not significantly affected by its presence. It is possible, however, that the influence of the complex extends into these adjacent regions, which would lead to an underestimation of the full remodelling effect. Nevertheless, we are confident in our findings for two main reasons. First, the method shows high reproducibility, producing consistent curvature estimates across multiple sperm cells (six cells) and three slices from each sperm cell (Table S1). Second, the results were validated using two separate computational techniques to measure changes in curvature—one based on the classic differential geometry formula to calculate the curvature from splines and another on a circle fitting into the arc cap—both yielding similar outcomes. The consistent results across samples and analytical methods suggest that our measurements provide a reliable quantification of the SUN5-Nesprin3 induced membrane remodelling.

## References

Abraham MJ, Murtola T, Schulz R, Páll S, Smith JC, Hess B, Lindahl E (2015) GROMACS: High performance molecular simulations through multi-level parallelism from laptops to supercomputers. SoftwareX 1–2: 19-25

Abramson J, Adler J, Dunger J, Evans R, Green T, Pritzel A, Ronneberger O, Willmore L, Ballard AJ, Bambrick J et al. (2024) Accurate structure prediction of biomolecular interactions with AlphaFold 3. Nature 630: 493–500

Barad BA, Medina M, Fuentes D, Wiseman RL, Grotjahn DA (2023) Quantifying organellar ultrastructure in cryo-electron tomography using a surface morphometrics pipeline. J Cell Biol 222

Berendsen HJC, Postma JPM, van Gunsteren WF, DiNola A, Haak JR (1984) Molecular dynamics with coupling to an external bath. The Journal of Chemical Physics 81: 3684–3690

Bernetti M, Bussi G (2020) Pressure control using stochastic cell rescaling. J Chem Phys 153: 114107

Best RB, Zhu X, Shim J, Lopes PE, Mittal J, Feig M, Mackerell AD, Jr. (2012) Optimization of the additive CHARMM all-atom protein force field targeting improved sampling of the backbone phi, psi and side-chain chi(1) and chi(2) dihedral angles. J Chem Theory Comput 8: 3257–3273

Bhaskara RM, Grumati P, Garcia-Pardo J, Kalayil S, Covarrubias-Pinto A, Chen W, Kudryashev M, Dikic I, Hummer G (2019) Curvature induction and membrane remodeling by FAM134B reticulon homology domain assist selective ER-phagy. Nat Commun 10: 2370

Burt A, Gaifas L, Dendooven T, Gutsche I (2021) A flexible framework for multi-particle refinement in cryo-electron tomography. PLoS Biol 19: e3001319

Castano-Diez D, Kudryashev M, Arheit M, Stahlberg H (2012) Dynamo: a flexible, user-friendly development tool for subtomogram averaging of cryo-EM data in high-performance computing environments. J Struct Biol 178: 139–151

Cavazza T, Takeda Y, Politi AZ, Aushev M, Aldag P, Baker C, Choudhary M, Bucevicius J, Lukinavicius G, Elder K et al. (2021) Parental genome unification is highly error-prone in mammalian embryos. Cell 184: 2860–2877 e2822

Cazin C, Boumerdassi Y, Martinez G, Fourati Ben Mustapha S, Whitfield M, Coutton C, Thierry-Mieg N, Di Pizio P, Rives N, Arnoult C et al. (2021) Identification and Characterization of the Most Common Genetic Variant Responsible for Acephalic Spermatozoa Syndrome in Men Originating from North Africa. Int J Mol Sci 22

Chaigne A, Campillo C, Voituriez R, Gov NS, Sykes C, Verlhac MH, Terret ME (2016) F-actin mechanics control spindle centring in the mouse zygote. Nat Commun 7: 10253

Cruz VE, Esra Demircioglu F, Schwartz TU (2020) Structural Analysis of Different LINC Complexes Reveals Distinct Binding Modes. J Mol Biol 432: 6028–6041

Darden T, York D, Pedersen L (1993) Particle mesh Ewald: An N⋅log(N) method for Ewald sums in large systems. The Journal of Chemical Physics 98: 10089–10092

Donahue RP (1972) Fertilization of the mouse oocyte: sequence and timing of nuclear progression to the two-cell stage. J Exp Zool 180: 305–318

Eisenstein F, Yanagisawa H, Kashihara H, Kikkawa M, Tsukita S, Danev R (2023) Parallel cryo electron tomography on in situ lamellae. Nat Methods 20: 131–138

Friedl P, Wolf K, Lammerding J (2011) Nuclear mechanics during cell migration. Curr Opin Cell Biol 23: 55–64

G. Bussi DD, M. Parrinello (2007) Canonical sampling through velocity rescaling. J Chem Phys 126

Galletta BJ, Ortega JM, Smith SL, Fagerstrom CJ, Fear JM, Mahadevaraju S, Oliver B, Rusan NM (2020) Sperm Head-Tail Linkage Requires Restriction of Pericentriolar Material to the Proximal Centriole End. Dev Cell 53: 86–101 e107

Giassetti MI, Miao D, Law NC, Oatley MJ, Park J, Robinson LD, Maddison LA, Bernhardt ML, Oatley JM (2023) ARRDC5 expression is conserved in mammalian testes and required for normal sperm morphogenesis. Nat Commun 14: 2111

Gob E, Schmitt J, Benavente R, Alsheimer M (2010) Mammalian sperm head formation involves different polarization of two novel LINC complexes. PLoS One 5: e12072

Gurusaran M, Davies OR (2021) A molecular mechanism for LINC complex branching by structurally diverse SUN-KASH 6:6 assemblies. Elife 10

Gurusaran M, Erlandsen BS, Davies OR (2024) The crystal structure of SUN1-KASH6 reveals an asymmetric LINC complex architecture compatible with nuclear membrane insertion. Commun Biol 7: 138

Hennen J, Saunders CA, Mueller JD, Luxton GWG (2018) Fluorescence fluctuation spectroscopy reveals differential SUN protein oligomerization in living cells. Mol Biol Cell 29: 1003–1011

Hess B, Bekker H, Berendsen HJC, Fraaije JGEM (1997) LINCS: A linear constraint solver for molecular simulations. Journal of Computational Chemistry 18: 1463–1472

Humphrey W, Dalke, A. & Schulten, K. (1996) VMD: Visual molecular dynamics. J Mol Graph 14: 33–38

J. David Pajerowski KND, Franklin L. Zhong, Paul J. Sammak, and Dennis E. Discher (2007) Physical plasticity of the nucleus in stemcell differentiation. PNAS 104: 15619–15624

J. R. Kremer DNM, J. R. McIntosh (1996) Computer Visualization of Three-Dimensional Image Data Using IMOD. Journal of Structural Biology 116: 71–76

Jahed Z, Fadavi D, Vu UT, Asgari E, Luxton GWG, Mofrad MRK (2018) Molecular Insights into the Mechanisms of SUN1 Oligomerization in the Nuclear Envelope. Biophys J 114: 1190–1203

Jahed Z, Soheilypour M, Peyro M, Mofrad MR (2016) The LINC and NPC relationship -it’s complicated! J Cell Sci 129: 3219–3229

Jorgensen WL, Chandrasekhar J, Madura JD, Impey RW, Klein ML (1983) Comparison of simple potential functions for simulating liquid water. The Journal of Chemical Physics 79: 926–935

Karl Salzwedel JTW, Eric Hunter (1999) A Conserved Tryptophan-Rich Motif in the Membrane-Proximal Region of the Human Immunodeficiency Virus Type 1 gp41 Ectodomain Is Important for Env-Mediated Fusion and Virus Infectivity. Journal of Virology 73: 2469–2480

Kidmose RT, Juhl J, Nissen P, Boesen T, Karlsen JL, Pedersen BP (2019) Namdinator - automatic molecular dynamics flexible fitting of structural models into cryo-EM and crystallography experimental maps. IUCrJ 6: 526–531

Klauda JB, Venable RM, Freites JA, O’Connor JW, Tobias DJ, Mondragon-Ramirez C, Vorobyov I, MacKerell AD, Jr., Pastor RW (2010) Update of the CHARMM all-atom additive force field for lipids: validation on six lipid types. J Phys Chem B 114: 7830–7843

Kmonickova V, Frolikova M, Steger K, Komrskova K (2020) The Role of the LINC Complex in Sperm Development and Function. Int J Mol Sci 21

Kucinska MK, Fedry J, Galli C, Morone D, Raimondi A, Solda T, Forster F, Molinari M (2023) TMX4-driven LINC complex disassembly and asymmetric autophagy of the nuclear envelope upon acute ER stress. Nat Commun 14: 3497

Kundu P, Naskar D, McKie SJ, Dass S, Kanjee U, Introini V, Ferreira MU, Cicuta P, Duraisingh M, Deane JE et al. (2023) The structure of a Plasmodium vivax Tryptophan Rich Antigen domain suggests a lipid binding function for a pan-Plasmodium multi-gene family. Nat Commun 14: 5703

Lamm L, Zufferey, S., Righetto, R.D., Wietrzynski, W., Yamauchi, K.A., Burt, A., Liu, Y., Zhang, H., Martinez-Sanchez, A., Ziegler, S., Isensee, F., Schnabel, J.A., Engel, B.D., and Peng, T (2024) MemBrain v2: an end-to-end tool for the analysis of membranes in cryo-electron tomography. BioRxiv

Liebschner D, Afonine PV, Baker ML, Bunkoczi G, Chen VB, Croll TI, Hintze B, Hung LW, Jain S, McCoy AJ et al. (2019) Macromolecular structure determination using X-rays, neutrons and electrons: recent developments in Phenix. Acta Crystallogr D Struct Biol 75: 861–877

Lipowsky R (1991) The conformation of membranes. Nature 349: 475–481

Lombardi ML, Jaalouk DE, Shanahan CM, Burke B, Roux KJ, Lammerding J (2011) The interaction between nesprins and sun proteins at the nuclear envelope is critical for force transmission between the nucleus and cytoskeleton. J Biol Chem 286: 26743–26753

Lomize AL, Todd SC, Pogozheva ID (2022) Spatial arrangement of proteins in planar and curved membranes by PPM 3.0. Protein Sci 31: 209–220

Lowe DG (2004) Distinctive Image Features from Scale-Invariant Keypoints. Int J Comput Vis 60: 91–110

Lu W, Gotzmann J, Sironi L, Jaeger VM, Schneider M, Luke Y, Uhlen M, Szigyarto CA, Brachner A, Ellenberg J et al. (2008) Sun1 forms immobile macromolecular assemblies at the nuclear envelope. Biochim Biophys Acta 1783: 2415–2426

Majumder S, Hsu YY, Moghimianavval H, Andreas M, Giessen TW, Luxton GWG, Liu AP (2022) In Vitro Synthesis and Reconstitution Using Mammalian Cell-Free Lysates Enables the Systematic Study of the Regulation of LINC Complex Assembly. Biochemistry 61: 1495–1507

Manfrevola F, Guillou F, Fasano S, Pierantoni R, Chianese R (2021) LINCking the Nuclear Envelope to Sperm Architecture. Genes (Basel) 12

Mazharimousavi SH, Forghani, S. D., & Abtahi, S. N. (2017) Generalized Monge gauge. International Journal of Geometric Methods in Modern Physics 14: 1750062

McGillivary RM, Starr DA, Luxton GWG (2023) Building and breaking mechanical bridges between the nucleus and cytoskeleton: Regulation of LINC complex assembly and disassembly. Curr Opin Cell Biol 85: 102260

Meszaros N, Cibulka J, Mendiburo MJ, Romanauska A, Schneider M, Kohler A (2015) Nuclear pore basket proteins are tethered to the nuclear envelope and can regulate membrane curvature. Dev Cell 33: 285–298

Michaud-Agrawal N, Denning EJ, Woolf TB, Beckstein O (2011) MDAnalysis: a toolkit for the analysis of molecular dynamics simulations. J Comput Chem 32: 2319–2327

Morgan JT, Pfeiffer ER, Thirkill TL, Kumar P, Peng G, Fridolfsson HN, Douglas GC, Starr DA, Barakat AI (2011) Nesprin-3 regulates endothelial cell morphology, perinuclear cytoskeletal architecture, and flow-induced polarization. Mol Biol Cell 22: 4324–4334

Pettersen EF, Goddard TD, Huang CC, Meng EC, Couch GS, Croll TI, Morris JH, Ferrin TE (2021) UCSF ChimeraX: Structure visualization for researchers, educators, and developers. Protein Sci 30: 70–82

Rothballer A, Kutay U (2013) The diverse functional LINCs of the nuclear envelope to the cytoskeleton and chromatin. Chromosoma 122: 415–429

Sanchez KM, Kang G, Wu B, Kim JE (2011) Tryptophan-lipid interactions in membrane protein folding probed by ultraviolet resonance Raman and fluorescence spectroscopy. Biophys J 100: 2121–2130

Santos Ád, Knowles O, Dendooven T, Hale T, Hale VL, Burt A, Kolata P, Cannone G, Bellini D, Barford D et al. (2024) Human spermatogenesis leads to a reduced nuclear pore structure and function *BioRxiv*

Scheffler K, Uraji J, Jentoft I, Cavazza T, Monnich E, Mogessie B, Schuh M (2021) Two mechanisms drive pronuclear migration in mouse zygotes. Nat Commun 12: 841

Schertel A, Snaidero N, Han HM, Ruhwedel T, Laue M, Grabenbauer M, Mobius W (2013) Cryo FIB-SEM: volume imaging of cellular ultrastructure in native frozen specimens. J Struct Biol 184: 355–360

Schindelin J, Arganda-Carreras I, Frise E, Kaynig V, Longair M, Pietzsch T, Preibisch S, Rueden C, Saalfeld S, Schmid B et al. (2012) Fiji: an open-source platform for biological-image analysis. Nat Methods 9: 676–682

Sha YW, Xu X, Ji ZY, Lin SB, Wang X, Qiu PP, Zhou Y, Mei LB, Su ZY, Li L et al. (2018) Genetic contribution of SUN5 mutations to acephalic spermatozoa in Fujian China. Gene 647: 221–225

Shang Y, Yan J, Tang W, Liu C, Xiao S, Guo Y, Yuan L, Chen L, Jiang H, Guo X et al. (2018) Mechanistic insights into acephalic spermatozoa syndrome-associated mutations in the human SUN5 gene. J Biol Chem 293: 2395–2407

Shang Y, Zhu F, Wang L, Ouyang YC, Dong MZ, Liu C, Zhao H, Cui X, Ma D, Zhang Z et al. (2017) Essential role for SUN5 in anchoring sperm head to the tail. Elife 6

Sosa BA, Kutay U, Schwartz TU (2013) Structural insights into LINC complexes. Curr Opin Struct Biol 23: 285–291

Sosa BA, Rothballer A, Kutay U, Schwartz TU (2012) LINC complexes form by binding of three KASH peptides to domain interfaces of trimeric SUN proteins. Cell 149: 1035–1047

Starr DA, Fridolfsson HN (2010) Interactions between nuclei and the cytoskeleton are mediated by SUN-KASH nuclear-envelope bridges. Annu Rev Cell Dev Biol 26: 421–444

Steigmann DJ (2017) The role of mechanics in the study of lipid bilayer. Srpinger *(Ed)* 577

Tegunov D, Cramer P (2019) Real-time cryo-electron microscopy data preprocessing with Warp. Nat Methods 16: 1146–1152

Tegunov D, Xue L, Dienemann C, Cramer P, Mahamid J (2021) Multi-particle cryo-EM refinement with M visualizes ribosome-antibiotic complex at 3.5 A in cells. Nat Methods 18: 186–193

Turgay Y, Ungricht R, Rothballer A, Kiss A, Csucs G, Horvath P, Kutay U (2010) A classical NLS and the SUN domain contribute to the targeting of SUN2 to the inner nuclear membrane. EMBO J 29: 2262–2275

Wagner FR, Watanabe R, Schampers R, Singh D, Persoon H, Schaffer M, Fruhstorfer P, Plitzko J, Villa E (2020) Preparing samples from whole cells using focused-ion-beam milling for cryo-electron tomography. Nat Protoc 15: 2041–2070

Wang N, Tytell, J. & Ingber, D. (2009) Mechanotransduction at a distance: mechanically coupling the extracellular matrix with the nucleus. Nat Rev Mol Cell Biol 10: 75–82

Wang W, Shi Z, Jiao S, Chen C, Wang H, Liu G, Wang Q, Zhao Y, Greene MI, Zhou Z (2012) Structural insights into SUN-KASH complexes across the nuclear envelope. Cell Res 22: 1440–1452

Wu B, Gao H, Liu C, Li W (2020) The coupling apparatus of the sperm head and taildagger. Biol Reprod 102: 988–998

Wu EL, Cheng X, Jo S, Rui H, Song KC, Davila-Contreras EM, Qi Y, Lee J, Monje-Galvan V, Venable RM et al. (2014) CHARMM-GUI Membrane Builder toward realistic biological membrane simulations. J Comput Chem 35: 1997–2004

Xian-Zhen Jiang M-GY, Li-Hua Huang, Chang-Qi Li, Xiao-Wei Xing (2011) SPAG4L, a Novel Nuclear Envelope Protein Involved in the Meiotic Stage of Spermatogenesis. DNA and Cell Biology 30: 875–882

Xiang M, Wang Y, Wang K, Kong S, Lu M, Zhang J, Duan Z, Zha X, Shi X, Wang F et al. (2022) Novel Mutation and Deletion in SUN5 Cause Male Infertility with Acephalic Spermatozoa Syndrome. Reprod Sci 29: 646–651

Yerima G, Domkam N, Ornowski J, Jahed Z, Mofrad MRK (2023) Force transmission and SUN-KASH higher-order assembly in the LINC complex models. Biophys J 122: 4582–4597

Zhang Y, Liu C, Wu B, Li L, Li W, Yuan L (2021a) The missing linker between SUN5 and PMFBP1 in sperm head-tail coupling apparatus. Nat Commun 12: 4926

Zhang Y, Yang L, Huang L, Liu G, Nie X, Zhang X, Xing X (2021b) SUN5 Interacting With Nesprin3 Plays an Essential Role in Sperm Head-to-Tail Linkage: Research on Sun5 Gene Knockout Mice. Front Cell Dev Biol 9: 684826

Zheng S, Wolff G, Greenan G, Chen Z, Faas FGA, Barcena M, Koster AJ, Cheng Y, Agard DA (2022) AreTomo: An integrated software package for automated marker-free, motion-corrected cryo-electron tomographic alignment and reconstruction. J Struct Biol X 6: 100068

Zhou Z, Du X, Cai Z, Song X, Zhang H, Mizuno T, Suzuki E, Yee MR, Berezov A, Murali R et al. (2012) Structure of Sad1-UNC84 homology (SUN) domain defines features of molecular bridge in nuclear envelope. J Biol Chem 287: 5317–5326

Zhu F, Liu C, Wang F, Yang X, Zhang J, Wu H, Zhang Z, He X, Zhang Z, Zhou P et al. (2018) Mutations in PMFBP1 Cause Acephalic Spermatozoa Syndrome. Am J Hum Genet 103: 188–199

Zhu F, Wang F, Yang X, Zhang J, Wu H, Zhang Z, Zhang Z, He X, Zhou P, Wei Z et al. (2016) Biallelic SUN5 Mutations Cause Autosomal-Recessive Acephalic Spermatozoa Syndrome. Am J Hum Genet 99: 942–949

Zimmerli CE, Allegretti M, Rantos V, Goetz SK, Obarska-Kosinska A, Zagoriy I, Halavatyi A, Hummer G, Mahamid J, Kosinski J et al. (2021) Nuclear pores dilate and constrict in cellulo. Science 374: eabd9776

Zuiderveld K (1994) Contrast limited adaptive histogram equalization. Graphics gems IV: 474–485

