## Supplementary material for "*In-cell* structure of a LINC complex reveals the molecular basis for membrane remodelling and head-to-tail coupling in sperm cells": Supp_Table_1

| | | *SUN5-containing region length ($\mu m$)* | *Curvature ($\mu m^{-1}$) Method 1* | *Curvature ($\mu m^{-1}$) Method 2* |
|---|---|---|---|---|
| *Sperm 1* | Slice 1 | 1.07 | 1.68 | 1.45 |
| | Slice 2 | 1.11 | 1.60 | 1.38 |
| | Slice 3 | 1.10 | 1.78 | 1.38 |
| *Sperm 2* | Slice 1 | 0.99 | 1.72 | 1.46 |
| | Slice 2 | 1.04 | 1.68 | 1.41 |
| | Slice 3 | 1.05 | 1.96 | 1.41 |
| *Sperm 3* | Slice 1 | 1.20 | 1.60 | 1.36 |
| | Slice 2 | 1.24 | 1.73 | 1.36 |
| | Slice 3 | 1.28 | 1.75 | 1.29 |
| *Sperm 4* | Slice 1 | 1.17 | 1.52 | 1.23 |
| | Slice 2 | 1.17 | 1.44 | 1.12 |
| | Slice 3 | 1.11 | 1.60 | 1.23 |
| *Sperm 5* | Slice 1 | 1.16 | 1.66 | 1.28 |
| | Slice 2 | 1.21 | 1.56 | 1.30 |
| | Slice 3 | 1.25 | 1.70 | 1.29 |
| *Sperm 6* | Slice 1 | 1.14 | 2.12 | 1.55 |
| | Slice 2 | 1.11 | 2.31 | 1.64 |
| | Slice 3 | 1.21 | 2.07 | 1.43 |
| *Mean* | | 1.14 | 1.75 | 1.37 |
| *Std* | | 0.08 | 0.22 | 0.12 |
