## Supplementary material for "*In-cell* structure of a LINC complex reveals the molecular basis for membrane remodelling and head-to-tail coupling in sperm cells": Supp_Table_2

| System | Description | Time (μs) | System Size (No. of atoms) | System size (nm) |
|---|---|---|---|---|
| SUN5_INM | Trimer of SUN5 inner membrane domain (55-126) + membrane. | 2.55 (0.85 × 3 replicas) | 291218 | 19.6 × 19.6 × 9.9 |
| SUN5_INM_AMPHI | Amphipathic helix of SUN5 (65-85) and membrane | 0.45 (0.15 × 3 replicas) | 68465 | 7.4 × 7.4 × 12.1 |
| SUN5_Nesprin3_WT | Trimer of WT SUN5-Nesprin3 complex (177-361 and 901-971) + membrane. | 2.1 (0.7 × 3 replicas) | 365105 | 17.8 × 17.6 × 15.0 |
| SUN5_trimer_membrane_hairpin | Trimer of SUN5 SUN domain (177-364) + membrane. KASH-lid loops in hairpin conformation | 3.0 (1.0 x 3 replicas) | 157505 | 11.1 x 11.1 x 13.7 |
| SUN5_trimer_membrane_nohairpin | Trimer of SUN5 SUN domain (177-364) + membrane. KASH-lid loops disordered | 3.0 (1.0 x 3 replicas) | 162894 | 11.1 x 11.1 x 14 |
| SUN5_trimer_membrane_solution model | Trimer of SUN5 SUN domain (177-364) + membrane. KASH-lid loops in folded conformation (~SUN2 crystal structure) | 0.52 (0.18 x 3 replicas) | 173078 | 12.0 x 12.0 x 11.7 |
| SUN5_trimer_solution | SUN5 trimer with KASH-lid loops in hairpin conformation in solution | 0.5 (0.5 x 1 replicas) | 110545 | 10.3 x 10.3 x 10.3 |
| SUN5_Nesprin3-trimer_solution | SUN5-Nesprin3 trimer (177-361 and 948-971) in solution | 0.5 (0.5 x 1 replicas) | 111859 | 10.3 x 10.3 x 10.3 |
| SUN5_Nesprin_FLHexamer_doublemembrane | SUN5-Nesprin3 Full-length hexamer modelled in double bilayer system | 0.6 (0.2 x 3 replicas) | 2836005 | 30.7 x 30.7 x 29.3 |
| SUN5_Nesprin_FLTrimer_doublemembrane | SUN5-Nesprin3 Full-length trimer modelled in double bilayer system | 1.5 (0.5 x 3 replicas) | 893882 | 17.1 x 17.1 x 29.6 |

| | | | | |
|---|---|---|---|---|
| SUN5_Nesprin3_Hexamer | Hexagonal cluster of SUN5-Nesprin3 complex (177-361 and 901-971) + membrane. | 2.28 (0.76 × 3 replicas) | 1984772 | 35.1 × 35.3 × 18.8 |
| SUN5_Nesprin3_NoTM_Hexamer | Hexagonal cluster of SUN5-Nesprin3 complex (177-361 and 948-971) without Nesprin3 transmembrane domain + membrane. | 2.7 (0.9 × 3 replicas) | 1456464 | 34.0 × 34.0 × 13.3 |
| SUN5_Hexamer | Hexagonal cluster of SUN5 (177-361) outer membrane domain + membrane. | 2.9 (0.96 × 3 replicas) | 1464576 | 34.0 × 34.0 × 13.3 |
| SUN5_Nesprin2_Hexamer | Hexagonal cluster of SUN5-Nesprin2 complex (48-71 + 177-361) + membrane. | 1.48 (0.7 × 1 replica + 0.39 × 2 replicas) | 1462631 | 32.4 × 32.4 × 13.8 |
| SUN5_Nesprin3_Hexamer | Hexagonal cluster of SUN5-Nesprin3 complex (177-361 and 901-971) + homogenous membrane (POPC). | 1.5 (0.5 × 3 replicas) | 1573465 | 35.1 × 35.1 × 14.7 |
| SUN5_Nesprin3_Trimer_Hexamer | Trimer of hexamer cluster of SUN5-Nesprin3 complex (177-361 and 901-971) + homogenous membrane (POPC). | 1.5 (0.5 × 3 replicas) | 2205521 | 40.0 × 40.0 × 14.8 |
| SUN5_Nesprin3_Hepta_Hexamer | Heptamer of hexamer cluster of SUN5-Nesprin3 complex (177-361 and 901-971) + homogenous membrane (POPC). | 1.5 (0.5 × 3 replicas) | 3488736 | 50.3 × 50.3 × 15.3 |
| Membrane | A heterogenous membrane | 0.5 (0.5 × 1 replicas) | 468510 | 31.4 × 31.4 × 9.1 |
